# Low Dose Radiation by Radiopharmaceutical Therapy Enhances GD2 *TRAC-*CAR T Cells Efficacy in Localized Neuroblastoma

**DOI:** 10.1101/2024.11.02.621668

**Authors:** Quaovi H. Sodji, Amanda Shea, Dan Cappabianca, Matthew H. Forsberg, Jens C. Eickhoff, Malick Bio Idrissou, Andy S. Ollendorff, Ohyun Kwon, Irene M. Ong, Reinier Hernandez, Jamey Weichert, Bryan P. Bednarz, Krishanu Saha, Paul M. Sondel, Christian M. Capitini, Zachary S. Morris

**Author notes:** **Corresponding author:** Quaovi H. Sodji, 600 Highland Ave, Madison, WI 53792, United States. Equal contribution (Christian M. Capitini, Zachary S. Morris).

## Abstract

**Background:** While chimeric antigen receptor (CAR) T cells have achieved significant success against hematological malignancies, efficacy against neuroblastoma has been limited. Virus-free CRISPR-edited GD2 *TRAC-*CAR T cells have been developed as a potential means of improving CAR T efficacy but are not curative. Radiopharmaceutical therapy (RPT) is a promising approach to enhance the effectiveness of immunotherapies, including immune checkpoint inhibitors. However, it remains unclear whether RPT can synergize with GD2 *TRAC-*CAR T cells to improve outcomes in neuroblastoma.

**Methods:** Dosimetry studies were conducted to measure the absorbed radiation dose delivered by lutetium-177 (^177^Lu) in both *in vitro* and *in vivo* models. Tumor-bearing mice were treated sequentially with low dose radiation by ^177^Lu-NM600, an alkylphosphocholine mimetic radiopharmaceutical agent, followed 9 days later by GD2 *TRAC-*CAR T cells generated in a virus-free manner by CRISPR/Cas9. Tumor burden was monitored through bioluminescence imaging and tumor size measurements. Mechanistic studies were performed using flow cytometry, multiplex assay and single-cell proteomic analysis.

**Results:** Low dose radiation delivered by ^177^Lu-NM600 synergized with GD2 *TRAC-*CAR T cells in a localized neuroblastoma model, resulting in complete tumor regression in all mice. The optimal combination was dependent on both the radiation dose and timing to minimize the negative impact of radiation on CAR T cell viability. Irradiation of neuroblastoma cells by low-dose RPT before GD2 *TRAC-*CAR T cells enhanced the release by CAR T cells of perforin, granzyme B and cytokines like TNF-α and IL-7 while abrogating TGF-β1 secretion. Additionally, low-dose RPT upregulated Fas on neuroblastoma cells, potentially enabling a CAR-independent killing.

**Conclusions:** This study demonstrates that low-dose RPT can enhance CAR T cell efficacy to treat a solid tumor. Findings suggest that optimization of radiation dose and timing may be needed for each patient and RPT to account for effects of varied tumor radiosensitivity and dosimetry.

**Graphical Abstract:** 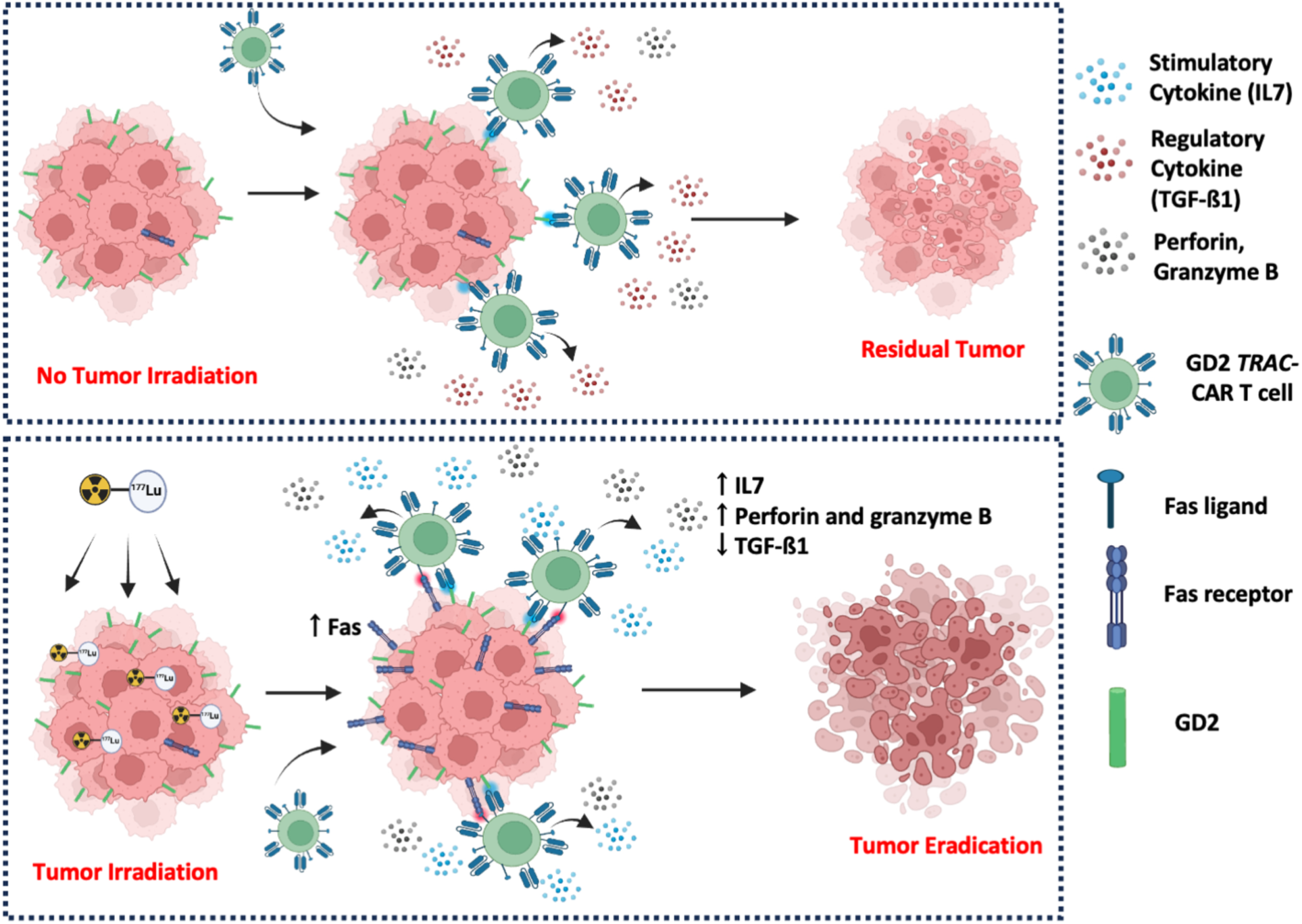

## BACKGROUND

Chimeric antigen receptor (CAR) T cell therapy has transformed the treatment of hematological malignancies. In patients with treatment-resistant CD19-expressing lymphoma and leukemia as well as B cell maturation antigen (BCMA)-expressing multiple myeloma, CAR T cells have achieved complete remission rate ranging from 54 to 90% ^1, 2, 3, 4^. However, this high efficacy has not been replicated in solid tumors, including neuroblastoma, the most common pediatric extracranial solid malignancy, which expresses high levels of the tumor-associated disialoganglioside GD2 ^5, 6, 7, 8^. This limited efficacy is in part due to limited CAR T cell infiltration into the tumor microenvironment (TME), exhaustion, poor persistence and antigen escape with little to no antigen expression by the tumor ^9^.

Development of virus-based third-generation CAR T cells targeting GD2 have recently demonstrated the highest clinical efficacy observed to date in treating relapsed or refractory high-risk neuroblastoma. In a cohort of treatment-resistant neuroblastoma patients, 33% achieved complete response (CR), 30% had partial response (PR) and 19% showed stable disease ^10^. However, 3-year event-free survival was modest at 36%, suggesting more advanced engineering of GD2 CAR T cells may be needed. Promising strategies such as CRISPR-Cas9 gene editing of CAR transgenes into the T cell receptor alpha constant (*TRAC*) gene have emerged. *TRAC-*CAR T cells lack endogenous T cell receptors (TCR) on the cell surface and demonstrate less tonic signaling with enrichment of stem cell memory phenotypes, greater potency and resistance to exhaustion compared to virally-manufactured CAR T cells ^11, 12, 13^.

Radiation promotes immune cell activation and migration into the TME by inducing immunogenic cell death ^14, 15^. Preclinical evidence suggests a potential synergy between radiation delivered as external beam radiation (EBRT) and CAR T cells against solid tumors. In a model of GD2-expressing glioblastoma, sub-lethal dose of focal or whole-body irradiation administered by EBRT combined with GD2 CAR T cells led to a durable complete tumor response. *In vivo* real-time imaging demonstrated that radiation enhanced CAR T cell infiltration and expansion within the TME ^16^. However, in cases of widespread metastasis where whole-body EBRT may be necessary to target multiple sites of disease to synergize CAR T cell therapy, concerns about acute and long-term toxicities arise. These include lymphopenia, endocrine dysfunctions, cardiovascular diseases, radiation pneumonitis, and infertility ^17^. Additional risks of secondary malignancies, growth failure and long-term neurocognitive deficits are particularly worrisome in pediatric patients ^18, 19^.

Radiopharmaceutical therapy (RPT) encompasses drugs that selectively deliver systemic radiation *in vivo* via a tumor-selective ligand labeled with a radionuclide with markedly less toxicity than whole-body EBRT ^20, 21^. Emerging evidence indicates that combining low-dose radiation from RPT with immune checkpoint inhibitors generates a robust antitumor response ^22^.

Herein, we aim to enhance the therapeutic efficacy of GD2 *TRAC-*CAR T cells against neuroblastoma using a low-dose radiation delivered by lutetium-177 (^177^Lu)-based RPT. We found that low-dose RPT enhances the efficacy of GD2 *TRAC*-CAR T cells, and both radiation dose and timing were critical for maximizing the antitumor response. Moreover, the enhanced cytotoxicity may be partly mediated by a CAR-independent killing mechanism driven by radiation-induced upregulation of the death receptor Fas on malignant cells.

## METHODS

### Cell lines

The human GD2+ tumor cell lines CHLA-20 (neuroblastoma) and M21 (melanoma), were generously provided by Dr. Mario Otto (University of Wisconsin-Madison) and Dr. Ralph Reisfeld (Scripps Research Institute), respectively. These cells were cultured in Dulbecco’s Modified Eagle Medium (DMEM) with high glucose (Gibco), supplemented with 10% fetal bovine serum (FBS) (Avantor) and 1% penicillin-streptomycin (Gibco). GD2+ SY5Y cells (neuroblastoma), donated by Dr. Crystal Mackall (Stanford University), were cultured in RPMI (Gibco) with 10% FBS and 1% penicillin-streptomycin. GD2- CCL-136 cells (rhabdomyosarcoma) were obtained from ATCC and grown in DMEM with 10% FBS and 1% penicillin-streptomycin. Tumor cell lines were lentivirally transduced with AkaLUC-GFP (VectorBuilder). All cells were maintained at 37 °C in a 5% CO_2_ atmosphere. Cell line authentication was performed using genomic short tandem repeat analysis (Idexx BioAnalytics) and morphological assessment according to ATCC guidelines. *Mycoplasma* contamination was routinely checked using the Mycoplasma Detection Kit MycoStrip™ (InvivoGen).

### Human T cell isolation and CAR T cell manufacturing

T cells were isolated from leukocyte reduction system (LRS) cones obtained from the Oklahoma Blood Institute (Oklahoma City, OK) using negative selection with the EasySep™ Human T Cell Isolation Kit (cat. no. 17951; STEMCELL Technologies). Alternatively, T cells were isolated from peripheral blood of healthy donors under a University of Wisconsin-Madison IRB-approved protocol (2017-1070) using the RosetteSep Human T Cell Enrichment Cocktail (cat. no. 15021, 15061; STEMCELL Technologies). The sequence for a third generation GD2-CD28-OX40-CD3ζ construct was generously provided by Malcolm Brenner (Baylor College of Medicine). GD2 *TRAC-* CAR T cells were manufactured by nucleofection of CRISPR-Cas9 into isolated T cells as previously described ^12^, and cultured in ImmunoCult-XF T Cell Expansion Medium (cat. no. 10981; STEMCELL Technologies) supplemented with 10 ng/mL IL-7 (cat. no. 207-IL-005/CF; BioTechne) and 10 ng/mL IL-15 (cat. no. 247-ILB-005/CF; BioTechne) and maintained at 37°C in 5% CO_2_ ^13^.

### *In vitro* dosimetry

*In vitro* dosimetry studies were initially conducted in a 6-well cell culture plate. The specific activities of ^177^Lu required to deliver 2 Gy to a cell monolayer at specific time points were estimated using the Geant4 Monte Carlo toolkit with an extension of RAPID ^23^. To validate these Monte Carlo predictions, serial dilutions of various ^177^Lu activity were prepared in cell culture medium and added to each well. Thermoluminescent dosimeters (TLDs) were placed at the center under each well and analyzed after 10 days by the University of Wisconsin-Madison Radiation Calibration Laboratory (Calibration Cert # 1664.01). A standard curve was generated and used to determine the ^177^Lu activity needed to achieve a given radiation dose, as previously described ^24^. For studies performed in a 96-well plate, a model of a flat-bottom 96-well plate was developed and used in Geant4 Monte Carlo to estimate the specific activities ^177^Lu needed to deliver 2 Gy at selected time points.

### *In vitro* GD2 *TRAC*-CAR T cell irradiation

GD2 *TRAC*-CAR T cells (10^6^ per well on a 6-well plate) were irradiated with 2 Gy by ^177^Lu diluted in 3 mL of ImmunoCult-XF T cell Expansion Medium (cat. no. 10981; STEMCELL Technologies) supplemented with 50 IU/mL of IL-2 over 3 days and then examined by flow cytometry for viability as previously described ^25^.

### *In vitro* irradiation of GD2 *TRAC*-CAR T cells in presence of tumor cells

A total of 100,000 GD2 *TRAC*-CAR T cells and 10,000 tumor cells were seeded in 100 µL of DMEM in a 96-well plate and irradiated with 2 Gy delivered over 3 days by ^177^Lu. After 3 days, the cells were harvested and washed 3 times with PBS. CAR T cell viability was determined by flow cytometry as previously reported ^25^.

### *In vitro* cytotoxicity activity of GD2 *TRAC-*CAR T cells against irradiated neuroblastoma cells

Two hundred and fifty thousand neuroblastoma cells were irradiated with 2 Gy over 6 days by ^177^Lu diluted in 3 mL of DMEM in a 6-well plate. The cells were harvested and washed 3 times with PBS. A trypan blue assay was performed to evaluate cell viability, and the viable irradiated neuroblastoma cells were co-cultured with GD2 *TRAC*-CAR T cells at an effector-to-target (E:T) ratio of 10:1 for 24 hours. The cytotoxic activity of the GD2 *TRAC*-CAR T cells against irradiated neuroblastoma cells was measured by flow cytometry ^25^.

### *In vitro* cytokine detection assays

Twenty-four hours after co-culturing GD2 *TRAC*-CAR T cells with irradiated neuroblastoma cells, 50 µL of supernatant was collected and the levels of selected cytokines (TNF-α, IFN-γ, IL-4 and IL- 13) were analyzed using the V-PLEX Proinflammatory Panel 1 Kit (Meso Scale Discovery) as previously described ^12^.

### Radiosynthesis of ^177^Lu-NM600

The radiolabeling of 2-(trimethylammonio)ethyl(18-(4-(2-(4,7,10-tris(carboxymethyl)-1,4,7,10- tetraazacyclododecan-1-yl)acetamido)phenyl)octadecyl) phosphate (NM600) with ¹⁷⁷Lu-chloride (SHINE Technologies) was performed as previously reported ^26^. Briefly, ¹⁷⁷Lu-chloride was mixed with 55–110 nmol of NM600 in 0.5 M NaOAc buffer (pH 5.5). After incubation at 90°C for 30 minutes, the ¹⁷⁷Lu-NM600 complex was purified using reverse-phase chromatography on Sep Pak C18 cartridges (Waters) and eluted in absolute ethanol under N₂ stream. ¹⁷⁷Lu-NM600 was reconstituted in saline with 0.4% v/v polysorbate 20. The radiosynthesis yield was > 95% with a radiochemical purity > 98% at apparent molar activity of 3.4 MBq/nmol.

### *In vivo* dosimetry

*In vivo* dosimetry of ^177^Lu-NM600 was estimated as previously reported ^23, 26^. The total activity in each organ was calculated by extrapolating %IA/g at given time points. A standard mouse model was used to convert organ-specific cumulative activity into absorbed dose per injected activity (Gy/MBq). Dose contributions from surrounding organs were included in calculations.

### Xenograft mouse model with flank tumor for GD2-*TRAC* CAR T cell treatment

All procedures were approved by the Institutional Animal Care and Use Committee at the University of Wisconsin-Madison (M005915; M005670; M006815). Six- to 8-week-old male and female NOD-*Rag1^null^IL2rg^null^* (NRG) mice (The Jackson Laboratory) were subcutaneously injected with 1 x 10^7^ luciferase-expressing CHLA-20 (AkaLUC-GFP CHLA-20) cells or SY5Y (AkaLUC-GFP SY5Y) or 1.5 × 10^6^ M21 (AkaLUC-GFP M21). Five days after injection, tumor implantation was confirmed by measuring tumor bioluminescence after the intraperitoneal (IP) injection of 150 mg/kg of D-luciferin using the *In Vivo* Imaging System (IVIS) (PerkinElmer). Randomization into the treatment groups was performed based on the BLI signal analyzed with Living Image Software (Revvity) and reported as total flux per second (photon/s). Mice receiving low dose RPT were administered ^177^Lu-NM600 on day 6 (post-tumor injection) and 3×10^6^ (CAR positive) GD2 *TRAC*- CAR T cells were administered on day 15. All mice were followed for toxicity, bioluminescence and overall survival. Disease burden was monitored *in vivo* through the tumor size and bioluminescence. Images were acquired weekly on the IVIS 15 minutes after IP injection of 150 mg/kg of D-luciferin. Tumor dimensions were measured bi-weekly with an electronic caliper and volumes were calculated as followed: (width)^2^ x (length/2). For survival studies, mice were euthanized when tumor volume reached 1500 mm^3^ or tumor ulceration was observed. Weight loss was also monitored for treatment-related toxicity like xenogeneic graft-versus-host-disease (GVHD). At select timepoints, tumors and spleens were harvested in FBS-free RPMI medium, mechanically dissociated and filtered through a 70 µm strainer (Corning). The resulting cell suspensions were centrifuged at 1,500 rpm for 5 minutes and the pellets were resuspended in ACK lysing buffer (Gibco) for 3 minutes. After the RBC lysis, 10 mL of PBS was added, and the samples were centrifuged again at 1,500 rpm for 5 minutes. The pellets were resuspended in PBS and analyzed by flow cytometry.

### Flow cytometry analysis

For *in vitro* analysis, cells were harvested and washed with PBS; samples obtained from *in vivo* studies were processed as described above. All specimens were resuspended into a single-cell solution in PBS as previously reported ^27^. Fc blocking with Human TruStain FcX™ (cat. no. 422302; Biolegend) and Live/Dead staining with Ghost Red Dye 780 (cat. no. 13-0865-T100; Tonbo Biosciences) were performed for 10 minutes at 4°C. The anti-GD2 scFV on the CAR was identified with 1A7 anti-14G2A antibody (cat. no. Ab02227-10.0; Absolute Antibody) labelled with APC via the APC Conjugation Kit Lightning-Link (cat. no. ab201807; Abcam). Other fluorophore-conjugated antibodies used include anti-GD2-PE-Dazzle594 (cat. no. 357320; Biolegend), anti-CD45-APC (cat. no. 982304; Biolegend) and anti-Fas-BV711 (cat. no. 305644; Biolegend). The fluorophore-conjugated antibodies were incubated for 20 minutes at 4°C and washed with 2% FBS in PBS. All analyses were performed using the Attune NxT Flow Cytometer (ThermoFisher) and data were analyzed with FlowJo version X (BD Biosciences).

### Single-cell multiplex secretome analysis

GD2 *TRAC*-CAR T cells were sorted by flow cytometry to enrich for CD8^+^ T cells. The sorted CD8^+^ GD2 *TRAC*-CAR T cells were co-cultured with previously irradiated (2 Gy delivered by ^177^Lu) or non-irradiated CHLA-20 cells at a E:T ratio of 1:2 for 20 hours. Stimulated CD8^+^ GD2 *TRAC*-CAR T cells were then carefully removed and stained with a cell membrane stain 405 (PhenomeX) and loaded onto the IsoCode chip. The single-cell secretome was measured during a 16-hour incubation. Secreted cytokines were identified using a panel of fluorescently labelled antibodies. The data generated were analyzed using IsoPeak software to evaluate the relative fluorescence intensity of the secreted cytokines, the polyfunctional strength, and to create a 3D UMAP of the CD8^+^ GD2 *TRAC*-CAR T cells ^28^.

### Statistical analysis

All data generated were analyzed using Prism v10 (GraphPad). Results are presented as mean ± standard deviation or standard error of the mean and compared between experimental conditions using two-sample *t*-tests or one-way analysis of variance (ANOVA), using Tukey’s Honestly Significant Difference (HSD) method for type I error control (<0.05) when conducting multiple pairwise comparisons. Survival outcomes were analyzed using the Kaplan-Meier method and compared between treatment groups using the log-rank test. All reported *p* values are two-sided and *p* <0.05 was used to define statistical significance.

## RESULTS

### Antigen stimulation of GD2 *TRAC-*CAR T cells during irradiation with ^177^Lu mitigates the deleterious effect of radiation on CAR T cell viability

GD2 *TRAC-*CAR T cells were engineered in a nonviral workflow using CRISPR/Cas9 ribonucleoproteins to knock-in a third generation GD2 CAR construct while knocking out the *TRAC* gene as previously described ^12^. The construct encoded a GD2-specific single-chain variable fragment 14g2a, a dual co-stimulatory domain with CD28 and OX40 and a CD3ζ cytotoxicity domain (Supplementary Fig. S1A). The timeline for GD2 *TRAC-*CAR T cell production was performed as previously described (Supplementary Fig. S1B) ^12^. We confirmed proteomic expression of the CAR and TCR knock out (KO) in T cells by flow cytometry and confirmed a robust TCR KO and GD2 CAR knock-in (Supplementary Fig. S1C). We previously observed *in vitro* that radiation delivered directly by ^177^Lu to GD2 *TRAC-*CAR T cells resulted in a dose-dependent CAR T cell death ^25^. This suggested that administering GD2 *TRAC-*CAR T cells after RPT, to minimize their exposure to radiation would be the optimal sequence for *in vivo* studies. Although this sequencing may reduce the irradiation of the GD2 *TRAC-*CAR T cells, complete avoidance of radiation is not possible. Therefore, we aimed to assess whether antigen stimulation during radiation exposure affects GD2 *TRAC-*CAR T cell viability. When radiation (2 Gy) was administered in the presence of GD2-expressing tumors such as CHLA-20 neuroblastoma or M21 melanoma (**Fig. 1A, 1B**, Supplementary Fig. S2), the viability of GD2 *TRAC-*CAR T cells is significantly improved. The presence of CHLA-20 and M21 cells led to a 2-fold and 3.5-fold increase in GD2 *TRAC-*CAR T cell viability, respectively (**Fig. 1C**; *p*< 0.0001). In contrast, no statistically significant increase in GD2 *TRAC-*CAR T cell viability was observed in the presence of the GD2-negative CCL-136 rhabdomyosarcoma (Supplementary Fig. S3B), (**Fig. 1B, 1C**). This enhanced viability of GD2 *TRAC-* CAR T cell viability in the presence of the GD2-expressing cell lines CHLA-20 and M21 was lost when antigen stimulation was delayed until after irradiation (Supplementary Fig. S3C).

**Figure 1:**
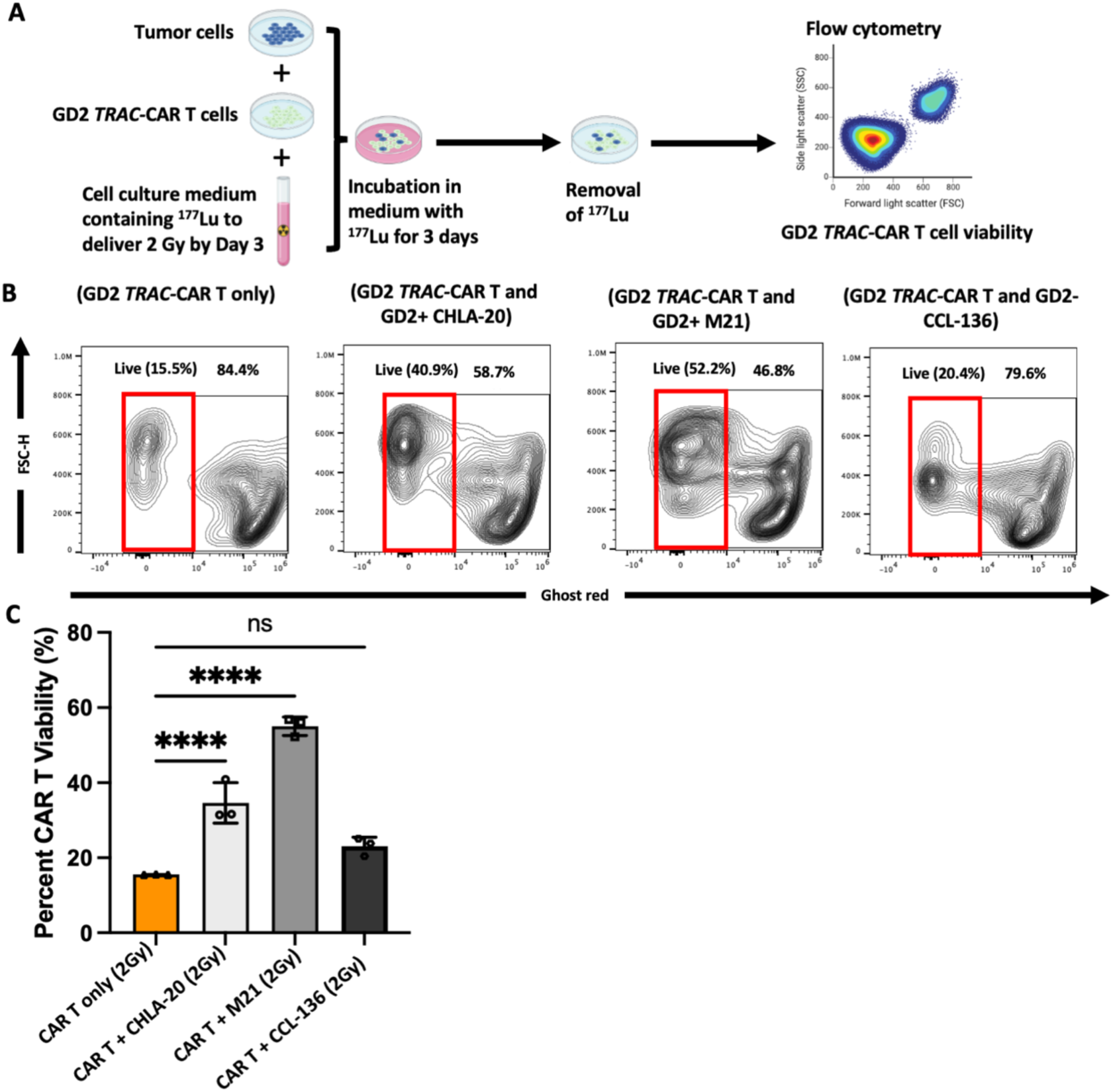
Antigen stimulation of GD2 *TRAC-*CAR T cells mitigates the deleterious effect of ^177^Lu radioisotope exposure on CAR T cell viability. **A)** CHLA-20 cells were co-cultured with GD2 *TRAC-* CAR T cells at an effector-to-target (E:T) ratio of 10:1 in medium containing ^177^Lu activity capable of delivering 2 Gy to all cells. After 3 days, ^177^Lu was removed and CAR T cell viability was assessed by flow cytometry. **B)** Representative contour plot illustrating the viability of irradiated (2 Gy) GD2 *TRAC-*CAR T cells (CD45^+^) co-cultured with GD2+ tumors (CHLA-20 and M21) and a GD2- tumor (CCL-136). **C)** Viability of GD2 *TRAC-*CAR T cells exposed to ^177^Lu radioisotopes after antigen dependent versus antigen independent stimulation. ****, *p* <0.0001. Error bars indicate SD. ns: not significant.

### Radiation delivered by ^177^Lu to neuroblastoma cells enhances their vulnerability to GD2 *TRAC-*CAR T cell mediated killing and stimulates cytokine secretion

We evaluated whether low-dose radiation by ^177^Lu sensitizes neuroblastoma cells to killing by GD2 *TRAC-*CAR T cells. To this end, CHLA-20 cells were irradiated with 2 Gy delivered by ^177^Lu over 6 days. After irradiation, the viable CHLA-20 cells, determined by a trypan blue assay, were co-cultured with GD2 *TRAC-*CAR T cells at a 10:1 effector to target (E:T) ratio for 24 hours (**Fig. 2A**). Compared to non-irradiated CHLA-20 cells, radiation alone (2 Gy) and GD2 *TRAC-*CAR T cells alone reduced tumor cell viability by 72.6% and 88% respectively. However, sequential tumor irradiation followed by GD2 *TRAC-*CAR T cells resulted in near-complete tumor cell eradication, with a 98% reduction in tumor cell viability (**Fig. 2B**; *p*< 0.0001). Additionally, prior irradiation of CHLA-20 cells led to a significant increase in TNF-α secretion by GD2 *TRAC-*CAR T cells compared to non-irradiated tumor cells (**Fig. 2C**, left; *p*= 0.01). No statistically significant differences were observed with secretion of other cytokines, including IFN*-*γ (**Fig. 2C**, right), IL-4 or IL-13 (**Fig. 2D**).

**Figure 2:**
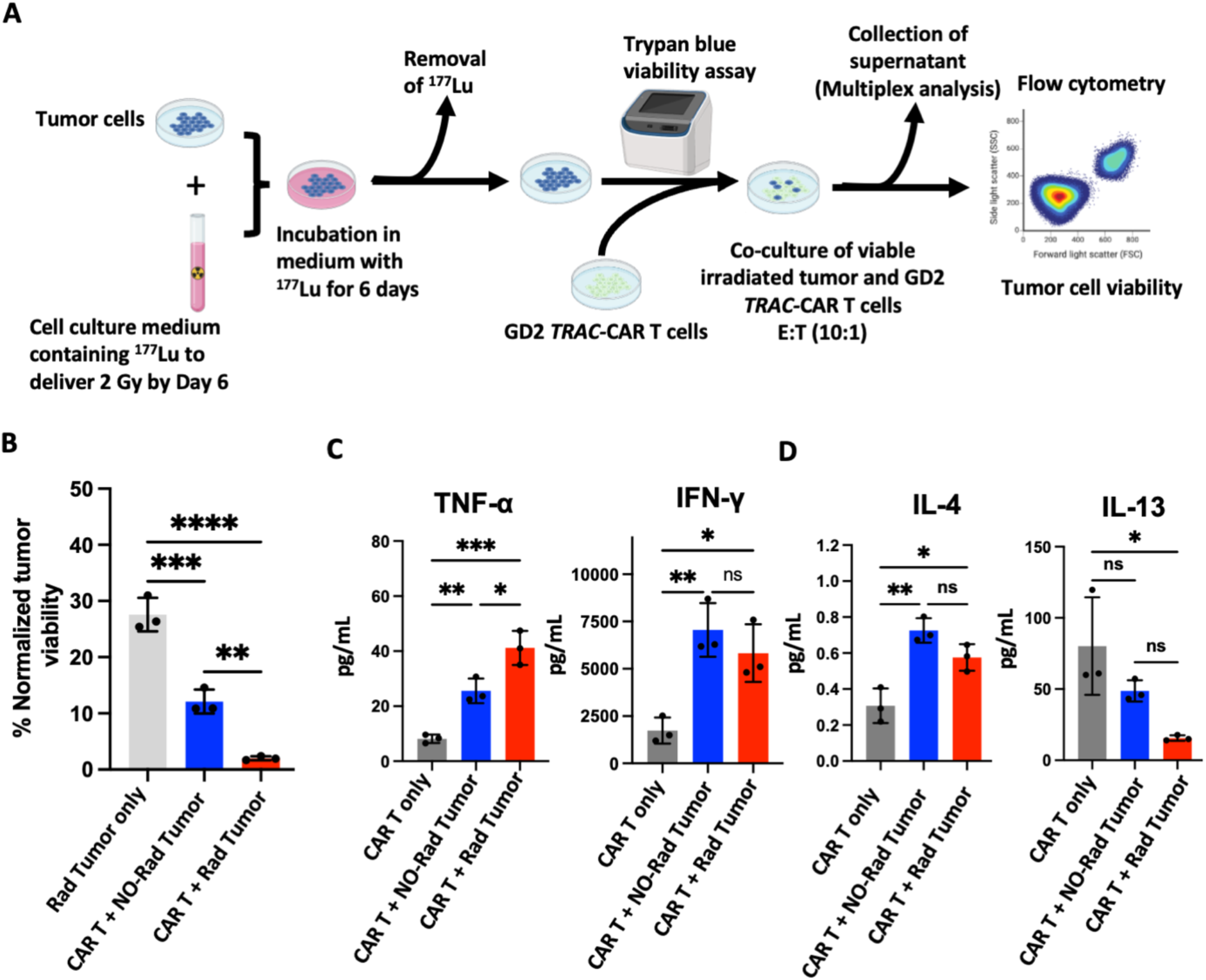
Radiation delivered by ^177^Lu to neuroblastoma cells enhances their vulnerability to GD2 *TRAC-*CAR T cell mediated killing and stimulates cytokine secretion. **A)** CHLA-20 cells were cultured in a medium containing ^177^Lu, delivering 2 Gy of radiation to tumor cells by Day 6. Following radiation delivery, ^177^Lu was removed and tumor viability was determined using a trypan blue assay. Viable CHLA-20 cells, irradiated and non-irradiated, were then co-cultured with GD2 *TRAC-*CAR T cells at a 10:1 E:T ratio for 24 hrs and tumor viability measured by flow cytometry. **B)** Viability of irradiated CHLA-20 cells after co-culture with GD2 *TRAC-*CAR T cells compared to radiation or *TRAC-*CAR T cells alone. **, *p*: 0.0027; ***, *p*: 0.0003; ****, *p*<0.0001. **C)** Secretion of T helper-1 (Th1) cytokines TNF-α and IFN-γ and **D)** Th2 cytokines IL-4 and IL-13 by GD2 *TRAC-*CAR T cells after their co-culture with irradiated CHLA-20 cells compared to non-irradiated CHLA-20 cells. TNF-α: *, *p*: 0.0126; **, *p*: 0.0076; ***, *p*: 0.0003. IFN-γ: *, *p*: 0.0176; **, *p*: 0.0051. IL-4: *, *p*: 0.0150; **, *p*: 0.0017. IL-13: *, *p*: 0.0186. Error bars indicate SD. ns: not significant.

### Assessing dosimetry and safety of ^177^Lu- NM600 in tumor xenograft mouse models

NM600 (**Fig. 3A**, top panel), is an alkylphosphocholine (APC) analog that selectively delivers radionuclides such as yttrium-90 (^90^Y) and ^177^Lu (^177^Lu-NM600; **Fig. 3A**, bottom panel) *in vivo* to a wide range of tumor types, including breast, prostate, lung and sarcoma ^26, 29^. Therefore, we investigated whether NM600 can deliver ^177^Lu in a neuroblastoma xenograft model in NRG mice. Longitudinal SPECT/CT studies were conducted after the intravenous (IV) injection of ^177^Lu- NM600 in NRG mice bearing CHLA-20 tumors (**Fig. 3B**). Representative maximum-intensity projections (MIP) of ^177^Lu-NM600 *in vivo* at various time points are shown in **Fig. 3C**, confirming the effective delivery of ^177^Lu by NM600 to the tumor. An *ex vivo* biodistribution study was performed 180 hours after the IV injection of ^177^Lu-NM600, revealing the accumulation of ^177^Lu- NM600 in the tumor and hepatic excretion consistent with previous reports (**Fig. 3D**) ^26^. Similarly, NM600 also delivers ^177^Lu in other xenograft models of GD2-expressing tumors including the melanoma M21 (Supplementary Fig. S4) and the neuroblastoma SY5Y (Supplementary Fig. S5A). Longitudinal SPECT/CT scans were used for dosimetry studies, which estimated an absorbed dose of 0.035 Gy/µCi for the CHLA-20 tumor. Dosimetry estimates of the absorbed doses for M21 and SY5Y tumors were 0.023 Gy/µCi and 0.0219 Gy/µCi respectively. Although hepatic excretion of ^177^Lu-NM600 raises potential concerns for hepatotoxicity, no changes were observed in liver function tests at the highest radiation dose (100 μCi) used for therapeutic studies (**Fig. 3E**). Moreover, this radiation dose did not adversely affect the differential blood counts or kidney functions (**Fig. 3E**), demonstrating the safety of low dose radiation delivered by the activities of ^177^Lu-NM600 administered in our study.

**Figure 3:**
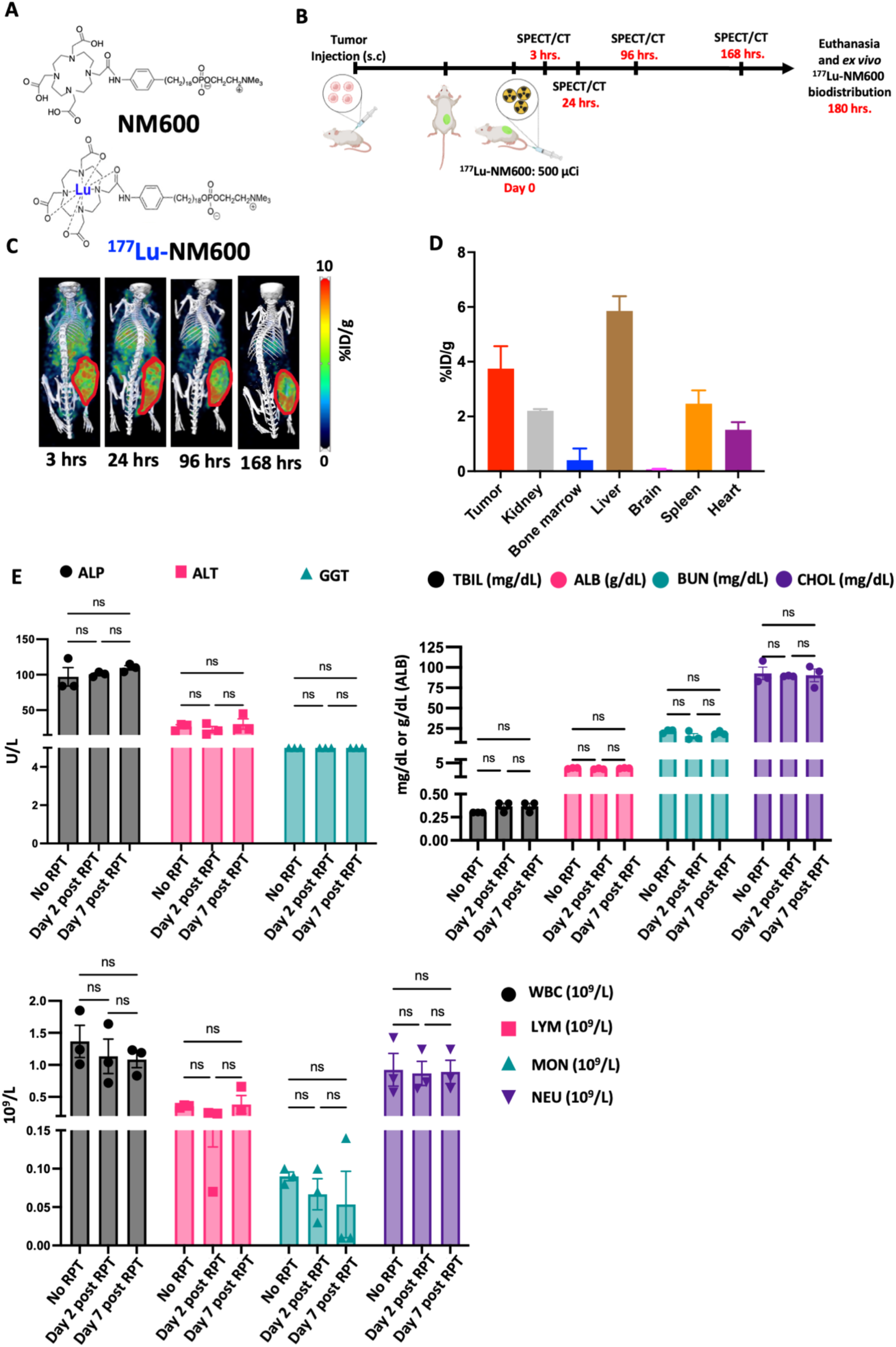
NM600 RPT delivers ^177^Lu to a human neuroblastoma xenograft mouse model. **A)** Flank CHLA-20 tumor-bearing NRG mice received ^177^Lu-NM600 (500 μCi) intravenously and serial SPECT/CT scans were performed after 3, 24, 96 and 168 hrs. After 180 hrs, the mice were euthanized and *ex vivo* biodistribution of ^177^Lu-NM600 was evaluated. **B)** Structure of unbound NM600 and NM600 chelating ^177^Lu (^177^Lu-NM600). **C)** Representative maximum intensity projection (MIP) SPECT/CT of ^177^Lu-NM600 in a CHLA-20 tumor-bearing NRG mouse. **D)** *Ex vivo* biodistribution of ^177^Lu-NM600 in the CHLA-20 xenograft model, 180 hrs after IV injection of ^177^Lu- NM600. **E)** The highest radiation dose of 100 μCi used for subsequent *in vivo* studies is not associated with end organ toxicities including hepatotoxicity (ALP, ALT, GGT, TBILI, ALB, CHOL), bone marrow toxicity (WBC, LYM, MON, NEU) and nephrotoxicity (BUN). ALK: Alkaline phosphatase; ALT: Alanine transaminase; GGT: Gamma-glutamyl transferase; TBIL: Total bilirubin; ALB: Albumin; BUN: Blood urea nitrogen; CHOL: Cholesterol; WBC: White blood cells; LYM: Lymphocytes; MON: monocytes; NEU: Neutrophils. ns: not statistically significant. Error bar: SD

### Low dose radiation delivered by ^177^Lu-NM600 enhances GD2 *TRAC-*CAR T cell therapy against a xenograft mouse model of localized neuroblastoma

Given the cooperative efficacy observed *in vitro* between radiation delivered by ^177^Lu and GD2 *TRAC-*CAR T cells, we evaluated this therapeutic effect in a human neuroblastoma flank xenograft model in NRG mice. We evaluated the *in vivo* efficacy of GD2 *TRAC-*CAR T cells when administered after two different low doses of radiation, 1.8 Gy and 3.6 Gy. After confirming the subcutaneous engraftment of luciferase-expressing CHLA-20 cells, tumor-bearing mice were randomized into four treatment groups: control (PBS), low dose radiation alone delivered by ^177^Lu-NM600, GD2 *TRAC-*CAR T cells alone or a combination group receiving low dose radiation followed by GD2 *TRAC-*CAR T cell infusion 9 days later (**Fig. 4A, 5A**). The delay between irradiation and GD2 *TRAC-* CAR T cell infusion was intended to minimize *TRAC-*CAR T cell exposure to radiation, allowing for the decay of ^177^Lu, which has a half-life of 6.6 days. Two methods—tumor volume and bioluminescence—were used to monitor therapeutic effects.

**Figure 4:**
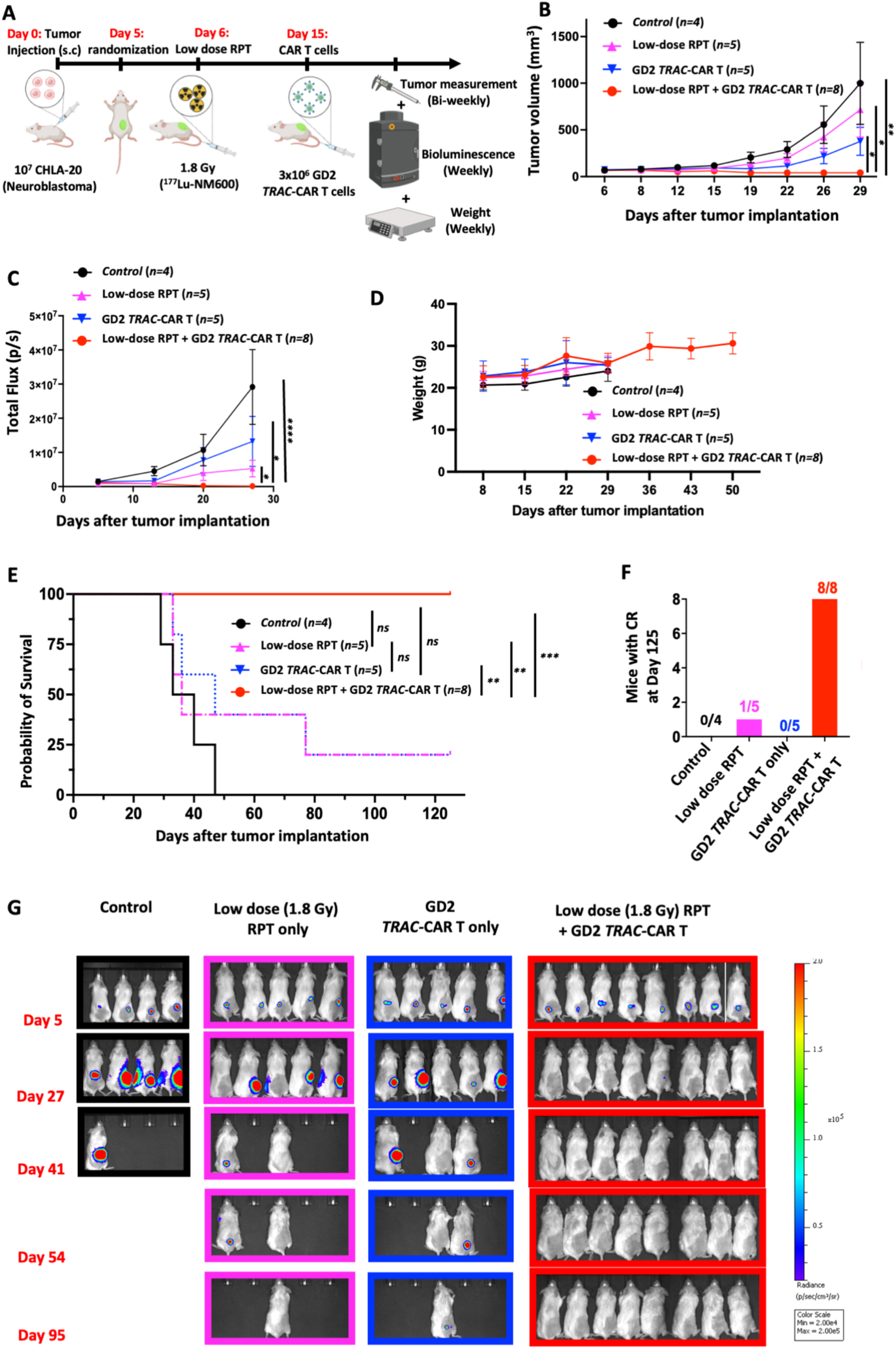
Low mean tumor dose of 1.8 Gy delivered *in vivo* by ^177^Lu-NM600 enhances GD2 *TRAC-* CAR T cell cytotoxicity against a localized human xenograft model of neuroblastoma and improves overall survival. **A)** Experimental scheme: 10^7^ CHLA-20 were subcutaneously injected on the right flank of NRG mice. After tumor implantation was confirmed by IVIS, tumor-bearing mice were randomized into 4 groups: control (PBS: n=4), low dose RPT alone (1.8 Gy mean tumor dose, n=5), GD2 *TRAC-*CAR T cells alone (3×10^6^ CAR+ cells) and low dose RPT followed by GD2 *TRAC-*CAR T cells (n=8) delivered 9 days post-RPT. Treatment effect was assessed by measuring tumor volume and bioluminescence. Treatment-related toxicity was measured by tracking weight loss and clinical assessments. **B)** Tumor volumes between the control group, monotherapy groups and combination therapy. Complete tumor regression by combination therapy was observed by Day 29 post-tumor injection (\**p*<0.012; \*\**p*=0.0087). **C)** Total flux (p/s) of tumors as measured by IVIS. Total flux was statistically lower in the combination group than in either monotherapy by Day 29 post-tumor injection (Two-way ANOVA with Dunnett’s multiple comparisons test; \**p*<0.01; \*\*\*\**p*<0.0001). **D)** Weights of each mouse was averaged for each group over time. **E)** Kaplan-Meier curves of the different treatment groups. All mice (8/8) in the combination group were alive 125 days post-tumor injection compared to 1 in 5 mice in either monotherapy group (\*\**p*=0.0019; \*\*\**p*=0.0002). **F)** Complete response in each treatment group at Day 125 post-tumor injection. All mice (8/8) had persistent complete response in the combination group compared 1 in 5 mice in the low dose RPT alone group and none in the control and anti-GD2 *TRAC-*CAR T cell group. **G)** Representative bioluminescence (radiance: p/sec/cm^2^/sr) images of all treatment groups.

**Figure 5:**
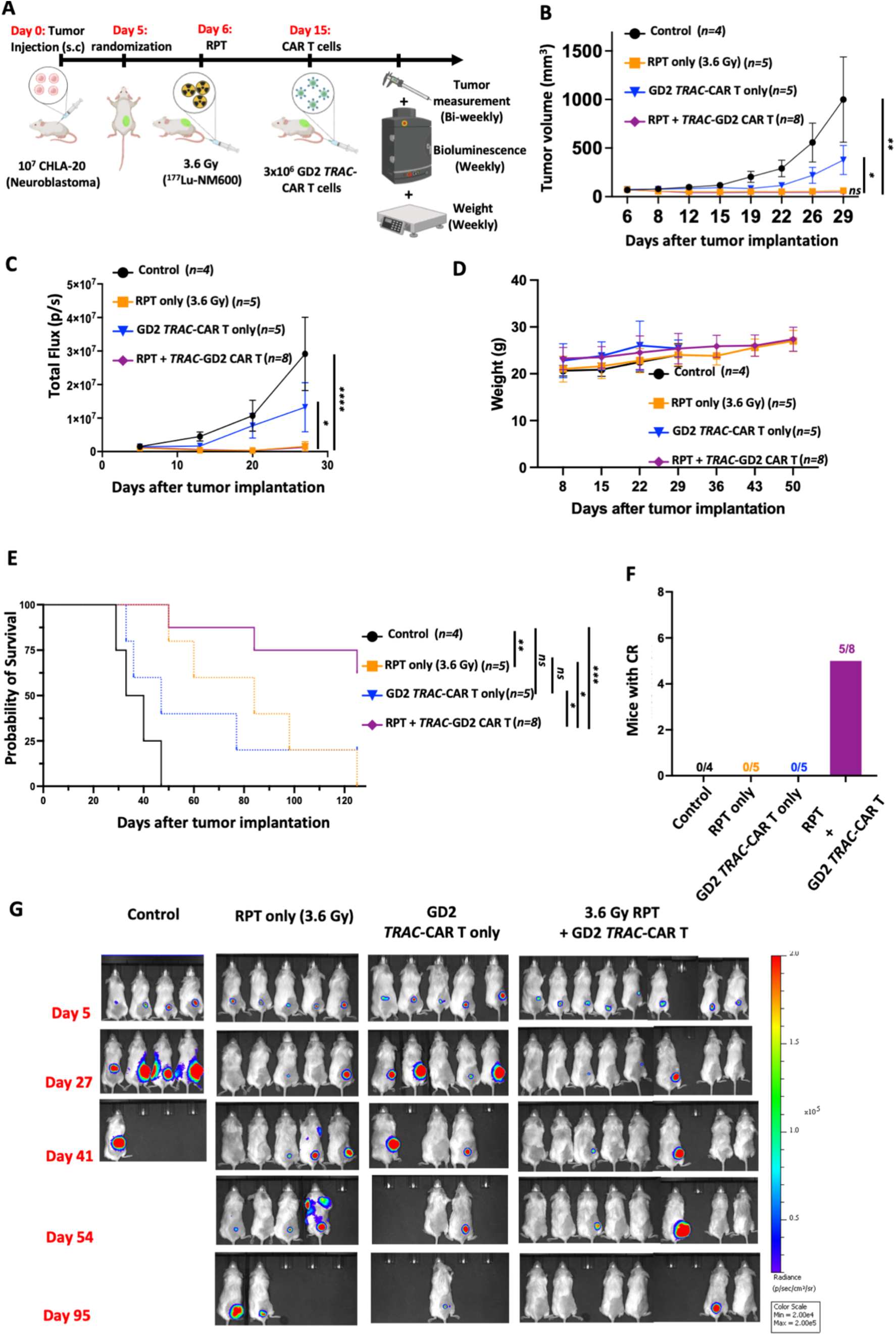
Low mean tumor dose radiation of 3.6 Gy delivered *in vivo* by ^177^Lu-NM600 improves efficacy of GD2 *TRAC-*CAR T cells against a localized human xenograft model of neuroblastoma. **A)** 10^7^ CHLA-20 were subcutaneously injected on the right flank of NRG mice. After tumor implantation was confirmed by IVIS, the tumor-bearing mice were randomized into 4 groups: control (PBS: n=4), 3.6 Gy mean tumor dose of RPT alone (n=5), GD2 *TRAC-*CAR T alone (3×10^6^ cells) and 3.6 Gy mean tumor dose from RPT followed by GD2 *TRAC-*CAR T cells (n=8) delivered 9 days post-RPT. Treatment effect was assessed by measuring tumor volume and bioluminescence. Treatment-related toxicity was measured by tracking weight loss and clinical assessment. **B)** Tumor volumes between the control group, GD2 *TRAC-*CAR T cells alone, 3.6 Gy RPT alone and the combination group (3.6 Gy RPT + GD2 *TRAC-*CAR T cells). Day 29 post-tumor injection, the combination led to significant tumor regression compared to the control and anti-GD2 *TRAC-*CAR T cell alone groups (\**p*=0.015; \*\**p*=0.009). **C)** Total flux (p/s) of tumors. Total flux was significantly lower in the combination group than in GD2 *TRAC-*CAR T cell alone and control group by Day 29 post-tumor injection (Two-way ANOVA with Dunnett’s multiple comparisons test; \**p*=0.023; \*\*\*\**p*<0.0001), but not statistical different from the 3.6 Gy monotherapy group. **D)** Weights were averaged in each group over time. **E)** Kaplan-Meier curves of the different treatment groups. 62.5% of mice (5/8) in the combination group were alive 125 days post-tumor injection compared to 0 in 5 mice in either monotherapy group (\**p*<0.035; \*\**p*=0.0027; *** *p*=0.0002). **F)** Complete response in each treatment group at Day 125 post-tumor injection. 5/8 mice had persistent complete response in the combination group compared none in all other groups. **G)** Representative bioluminescence (radiance: p/sec/cm^2^/sr) images of all treatment groups.

With a mean tumor dose from RPT of 1.8 Gy, combination RPT + GD2 CAR-*TRAC* therapy produced a robust treatment response, as compared to all control groups, leading to complete tumor regression in all mice by 29 days after tumor cell implantation (**Fig. 4B**, 4C). No treatment-related toxicity was observed, mice weights remained similar in all 4 groups (**Fig. 4D**), and there were no visible signs of xenogeneic GVHD in the groups receiving GD2 *TRAC-*CAR T cells, consistent with previous reports ^12, 30^. To assess the durability of response, mice were monitored for up to 125 days after tumor implantation. All mice (8 out of 8) in the combination group were alive at day 125, compared to 1 out of 5 in each monotherapy group and no mice in the untreated control group (**Fig. 4E**). Among the survivors, all mice receiving combination RPT + GD2 CAR-*TRAC* achieved a persistent CR, while only 1 CR was observed in the low dose RPT monotherapy group and no CRs were observed in the GD2 *TRAC-*CAR T cell monotherapy group (**Fig. 4F**).

Initially, the 3.6 Gy mean tumor dose RPT monotherapy group exhibited a therapeutic response similar to that of combination RPT with GD2 *TRAC-*CAR T cells (**Fig. 5B**, 5C). However, with extended follow up, combination therapy demonstrated a significantly better outcome compared with RPT monotherapy (**Fig. 5E**), with 5 out of 8 mice receiving the combination therapy showing a persistent CR up to 125 days post tumor implantation versus none of the mice receiving either monotherapy (**Fig. 5F**). The complete and persistent response rate for the combination of 3.6 Gy RPT and GD2 *TRAC*-CAR T cells was 62.5% (**Fig. 5F**) compared to 100% with the combination of 1.8 Gy RPT and GD2 *TRAC-*CAR T cells (**Fig. 4F**).

Similarly, the combination of low-dose RPT delivered by ^177^Lu-NM600 and GD2 *TRAC-*CAR T cells demonstrated an antitumor activity against a second GD2+ neuroblastoma xenograft model with SY5Y cells (Supplementary Fig. S5). This therapeutic combination also showed efficacy in targeting another tumor histology, the GD2+ melanoma M21, though with less potency than when tested against neuroblastoma (Supplementary Fig. S6).

### *In vivo* tumor irradiation with 1.8 Gy enhances GD2 *TRAC-*CAR T cells infiltration in the tumor microenvironment of neuroblastoma

Given that 1.8 Gy mean tumor dose from RPT demonstrated better efficacy when combined with GD2 *TRAC-*CAR T cells compared to 3.6 Gy, we evaluated the impact of low dose radiation from ^177^Lu-NM600 on the infiltration of GD2 *TRAC-*CAR T cells in the TME. To accomplish this, CHLA-20 tumor-bearing mice were treated with either GD2 *TRAC-*CAR T cells alone or in combination with low dose RPT (delivering 1.8 Gy mean tumor dose) administered before CAR T cell infusion, as previously done. Five days after the *TRAC-*CAR T cell infusion, tumors and spleens were harvested to assess the presence of GD2 *TRAC-*CAR T cells (**Fig. 6A**, Supplementary Fig. S7). Our results showed that prior irradiation led to nearly a 2-fold increase in GD2 *TRAC-*CAR T cells in the TME (**Fig. 6B**; *p*: 0.0052). However, no statistically significant difference was observed in the spleens of irradiated versus non-irradiated mice (**Fig. 6C**).

**Figure 6:**
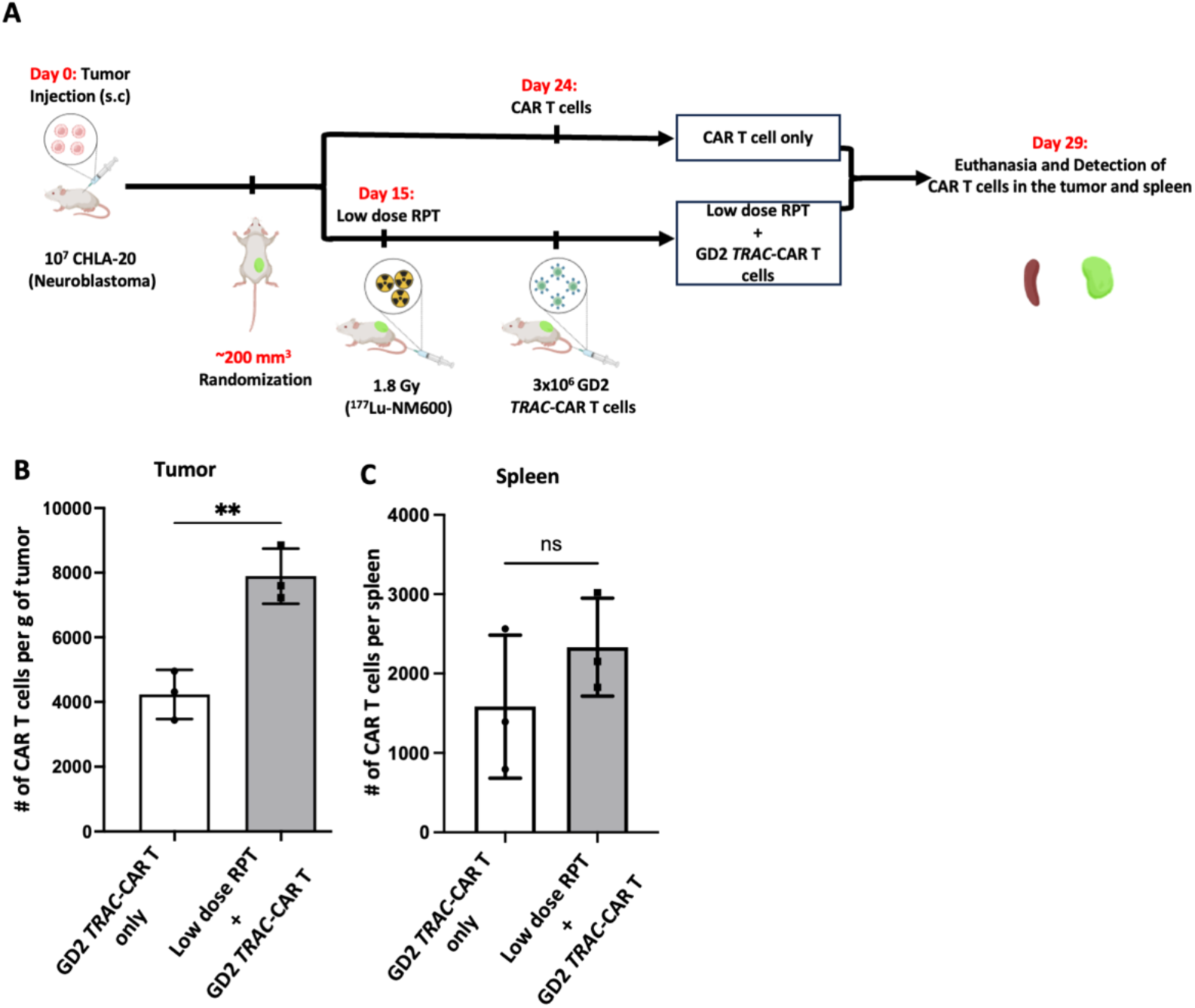
*In vivo* tumor irradiation with 1.8 Gy mean tumor dose enhances GD2 *TRAC-*CAR T cells infiltration in the tumor microenvironment of human neuroblastoma xenograft models. **A)** NRG mice bearing large CHLA-20 tumor of ∼200 mm^3^ were treated either with GD2 *TRAC-*CAR T cells or with the combination of 1.8 Gy mean tumor dose from RPT followed by CAR T cell infusion delivered 9 days later. Five days after CAR T cell infusion, the mice were euthanized and the number (#) of GD2 *TRAC-*CAR T cells per **B)** gram of tumor and **C)** in the whole spleen was analyzed by flow cytometry. 1.8 Gy mean tumor dose from RPT yielded ∼ a 2-fold increase infiltration of GD2 *TRAC-*CAR T cells in the TME compared to administering GD2 *TRAC-*CAR T cells alone (\*\**p*=0.0052). No statistical difference in GD2 *TRAC-*CAR T cells was noted in the spleen with either treatment. ns = not significant

### Irradiation of neuroblastoma enhances the polyfunctionality of CD8^+^ GD2 *TRAC-*CAR T cells

To assess the functional changes in CD8^+^ GD2 *TRAC-*CAR T cells induced by radiation from ^177^Lu on neuroblastoma, we analyzed the single-cell secretome of CD8^+^ GD2 *TRAC-*CAR T cells after their co-culture (stimulation) with either irradiated (2 Gy) or non-irradiated CHLA-20 neuroblastoma cells using the IsoCode chip (**Fig. 7A**) ^31^. Irradiation of neuroblastoma significantly increased the secretion by CD8^+^ GD2 *TRAC-*CAR T cells of granzyme B (GZMB) and perforin (PRF), while reducing the secretion of inflammatory proteins monocyte chemoattractant protein-1 (MCP-1) and MCP-4 (**Fig. 7B**). The secretion of chemoattractive cytokines macrophage inflammatory protein 1β (MIP-1β) and RANTES was slightly higher with non-irradiated tumor cells (**Fig. 7B**). The co-culturing of CD8^+^ GD2 *TRAC-*CAR T cells with irradiated tumor cells led to a significant increase in IL-7 secretion by the *TRAC-*CAR T cells, compared to those co-cultured with non-irradiated tumor cells. Furthermore, the tumor cell irradiation abolished the secretion of the regulatory/inhibitory cytokine TGF-β1 by CD8^+^ GD2 *TRAC-*CAR T cells (**Fig. 7B**).

**Figure 7:**
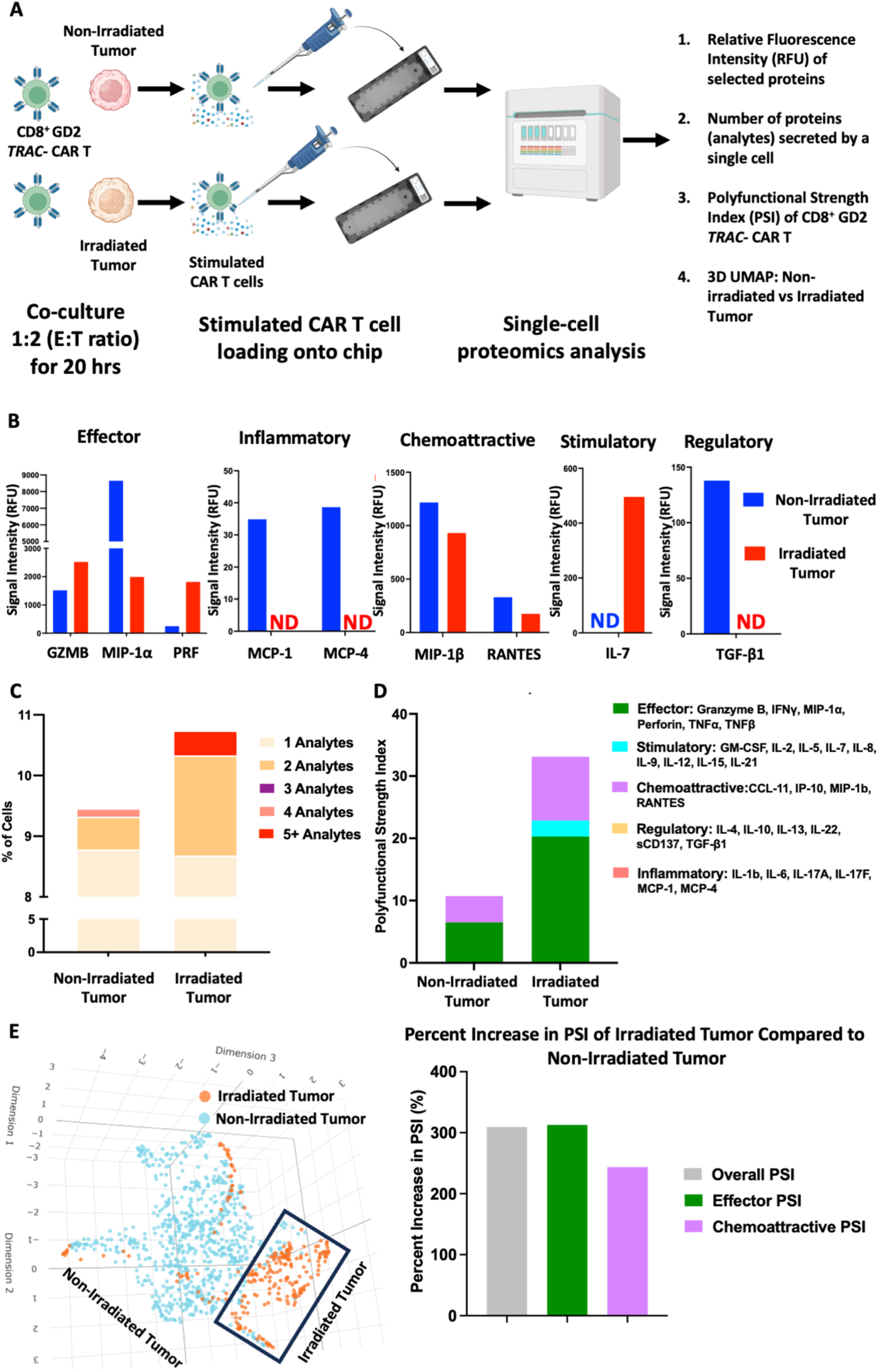
Irradiation of neuroblastoma induces a polyfunctional phenotype in CD8^+^ GD2 *TRAC-* CAR T cells. **A)** CD8^+^ GD2 *TRAC-*CAR T cells were co-cultured with irradiated or non-irradiated CHLA-20 tumor cells at an E:T ratio of 1:2 for 20 hrs. The CD8^+^ GD2 *TRAC-*CAR T cells were stained and loaded onto an IsoCode chip and the single-cell secretome was measured for 16 hrs. **B)** Relative fluorescence intensity (RFU) of selected cytokines/chemokines, categorized as effector, inflammatory, chemoattractive, stimulatory or regulatory, were measured. **C)** Number of individual proteins or analytes secreted by a single CAR T cell. **D)** Polyfunctional strength index (PSI) of CD8^+^ GD2 *TRAC-*CAR T cells when co-cultured with irradiated tumor cells versus non-irradiated tumor cells. **E)** 3D UMAP visualization of CD8^+^ GD2 *TRAC-*CAR T cells under different co-culture conditions. CD8^+^ GD2 *TRAC-*CAR T cells co-cultured with irradiated tumor display a distinctive and characteristic proteomic phenotype. ND: Not Detected.

Although the percentage of CD8^+^ GD2 *TRAC-*CAR T cells secreting a single cytokine was similar in both conditions, tumor cell irradiation led to a substantial increase in the population of CD8^+^ GD2 *TRAC-*CAR T cells secreting two or more cytokines (**Fig. 7C**). Notably, CD8^+^ GD2 *TRAC-*CAR T cells secreting five or more cytokines were detected following tumor irradiation (**Fig. 7C**). Compared to non-irradiated tumor cells, radiation delivered by ^177^Lu resulted in a 3-fold increase in the overall polyfunctional strength index (PSI) of CD8^+^ GD2 *TRAC-*CAR T cells. This increase was primarily driven by higher effector and chemoattractive PSIs (**Fig. 7D**). Moreover, a stimulatory PSI component, absent with non-irradiated tumor contributed to the overall PSI increase detected after tumor irradiation. Ultimately, the single-cell secretome analysis revealed that the irradiation of neuroblastoma altered the cytokine secretion profile by CD8^+^ GD2 *TRAC-*CAR T cells, leading to a distinct cluster of CD8^+^ GD2 *TRAC-*CAR T cells on a 3D UMAP, compared to non-irradiated neuroblastoma cells (**Fig. 7E**).

### Low-dose radiation from ^177^Lu increases the expression of the death receptor Fas on neuroblastoma cells, potentially mediating a CAR-independent killing mechanism by GD2 *TRAC*-CAR T cells

Radiation delivered by ^177^Lu to neuroblastoma cells enhances their sensitivity to GD2 *TRAC*-CAR T cell killing by boosting two known mechanisms of CAR T cell cytotoxicity: the release of cytokines such as TNF-α (**Fig. 2)** and cytotoxic effector proteins such as perforin and granzyme (**Fig. 7**) ^32^. We evaluated whether this radiation might also impact the death receptor killing pathway mediated by the interaction of Fas on tumor cells with its ligand FasL, a pathway known to trigger a CAR-independent killing by CAR T cells ^33^. To investigate this, CHLA-20 neuroblastoma cells were irradiated with a low dose of radiation (2 Gy), and their expression of Fas was assessed (**Fig. 8A**; Supplementary Fig. S8). We observed an approximately 15-fold increase in Fas expression following radiation (**Fig. 8B**; *p* <0.0001), along with a significant rise in median fluorescence intensity (MFI) (**Fig. 8C**; *p*: 0.0005). *In vivo* evaluation of the CAR-independent killing mechanism (Supplementary Fig. S9A) revealed that the combination of low-dose radiation (1.8 Gy) and mock (*mCherry) TRAC-*CAR T cells (Supplementary Fig. S9B), was ineffective with disease ultimately recurring in all mice (Supplementary Fig. S9C).

**Figure 8:**
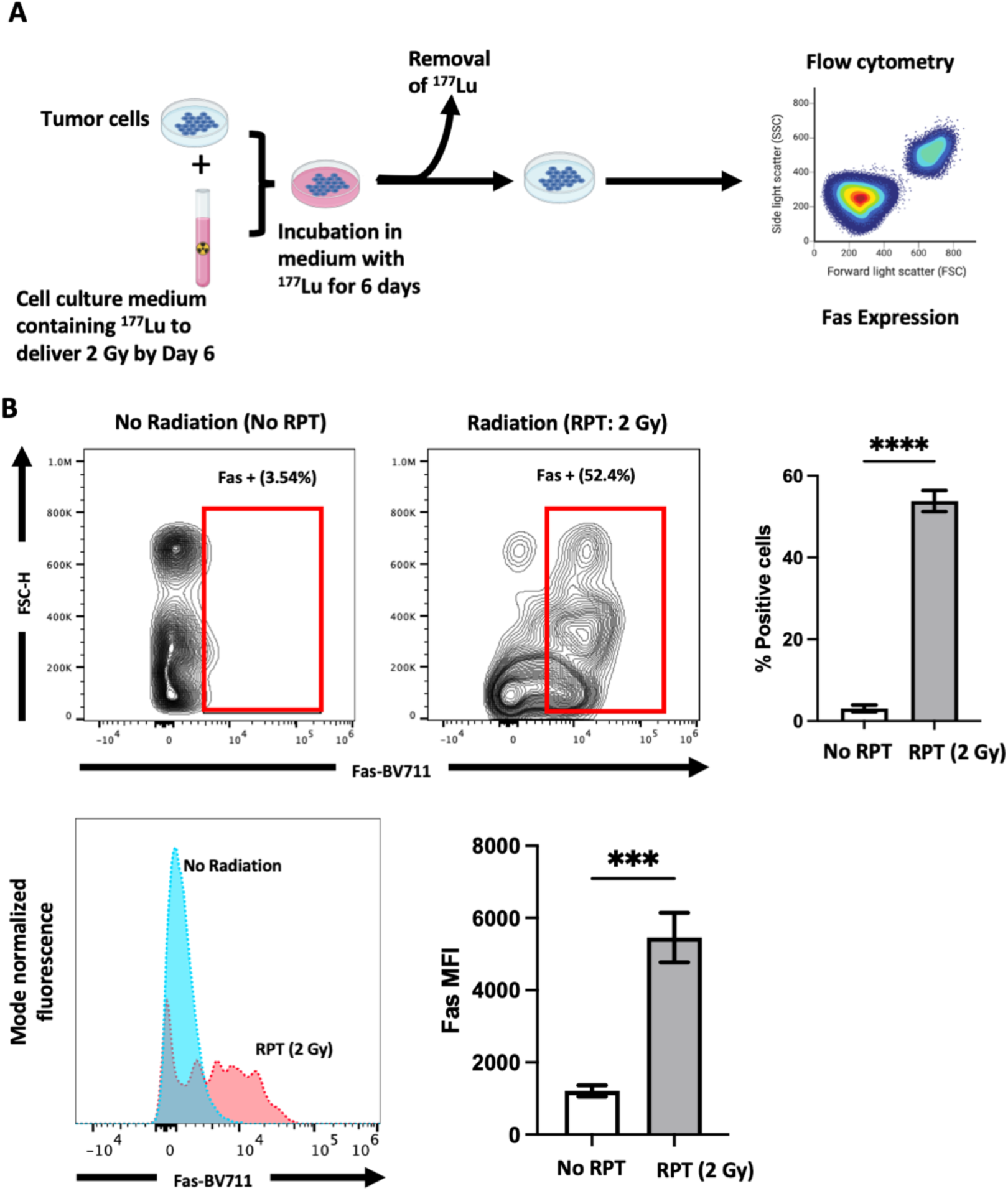
Irradiation of GD2+ neuroblastoma cells by ^177^Lu upregulates Fas expression. **A)** CHLA- 20 cells were incubated with ^177^Lu sufficient to deliver 2 Gy by day 6. Fas expression on tumor cells was evaluated by flow cytometry. **B)** Fas expression on irradiated neuroblastoma cells was significantly higher than that on non-irradiated neuroblastoma cells. ****, *p*< 0.0001. **C)** Mean fluorescence intensity (MFI) of Fas on irradiated neuroblastoma cells was also much higher compared to non-irradiated neuroblastoma cells. ***, *p*=0.0005

## DISCUSSION

Our study demonstrates the preclinical feasibility, safety and efficacy of combining low dose radiation delivered by RPT and GD2 *TRAC-*CAR T cell therapy against GD2+ solid tumors like neuroblastoma and melanoma. This strategy builds on previous strategies utilized to enhance CAR T cell therapy against neuroblastoma ^10, 12, 34^. In addition to enhancing GD2 *TRAC-*CAR T cell viability, cytotoxicity and cytokine secretion *in vitro*, complete responses and improved survival with high infiltration in the TME and no GVHD or end-organ toxicity was observed after testing 2 different low doses of RPT *in vivo*. Lower doses of RPT like 1.8 Gy mean tumor dose appeared to be more effective than 3.6 Gy mean tumor dose in enhancing survival and complete responses by GD2 *TRAC*-CAR T cells. Single-cell proteomics of CAR T cells showed enhanced secretion of multiple cytokines after exposure to irradiated tumors, especially within effector pathways. Low-dose radiation delivered by RPT may also promote a CAR-independent killing pathway mediated by Fas.

Radiation delivered by EBRT has been shown to potentiate CAR T cell cytotoxicity against various solid tumors ^16, 35, 36^. For instance, in a xenograft model of non-small cell lung cancer, tumor-targeted low-dose EBRT delivered before mesothelin-targeted CAR T cells enhanced their efficacy^37^. This low-dose EBRT promoted chemokine secretion by the tumor, leading to increased infiltration of CAR T cells with upregulation of corresponding chemokine receptors CXCR3, CXCR6, CCR2, CCR5 and CX3CR1 ^37^. While low-dose EBRT potentiates CAR T cell therapy against localized solid tumors, its use in metastatic settings may be limited by increased toxicity from larger radiation fields. In such instances, RPT may offer advantages such as the ability to target multiple sites of diseases, including radiographically occult lesions, with potentially less toxicity than EBRT^21^. However, RPT poses its own challenges due to the persistence of the radionuclide and ongoing radioactivity decay *in vivo*, potentially affecting adoptively transferred CAR T cells. A study by Awuah et al. demonstrated an improved antitumor activity against multiple myeloma when CS1- CAR T cells were administered 21 days after RPT (^225^Ac-DOTA-daratumumab) compared to 14 or 28 days ^38^. At 14 days post-RPT, residual radiation originating from the remaining RPT can still be detrimental to CAR T cell viability, while at 28 days, increase tumor burden can reduce CAR T cell efficacy. This underscores the importance of identifying optimal timing, order and dose for maximizing anti-tumor responses from combination RPT and CAR T cell therapy ^38^.

Yang and colleagues explored the combination of CAR T cells targeting intracellular adhesion molecule 1 (ICAM-1) or epithelial cellular adhesion molecule (EpCAM) with ^177^Lu-DOTATATE in xenograft models of gastric cancer ^39^. In their study, the CAR T cells were administered prior to RPT, specifically 28 and 16 days before receiving 7.4 MBq (2.6±1.17 Gy to tumor) or 37 MBq (6±1.35 Gy to tumor), respectively. Although the combination therapy delayed tumor growth compared to either monotherapy, the sequencing of the therapies implied that CAR T cells within the TME at the time of RPT were exposed to potentially harmful radiation, compromising their viability and subsequent cytotoxic activity ^39^. However, the exact mechanism behind the anti-tumor effects they observed remains unclear.

Minimizing radiation exposure to CAR T cells is paramount to a successful combination regimen. In previous work, we demonstrated that GD2 *TRAC-*CAR T cells exposed to 1 Gy of radiation delivered by ^177^Lu retained 87% of their viability *in vit*ro compared to non-irradiated GD2 *TRAC-*CAR T cells ^25^. This finding informed the timing of RPT and GD2 *TRAC-*CAR T cell administration during this current study. Given the 6.6-day half-life of ^177^Lu, when 1.8 Gy mean tumor dose from RPT (^177^Lu-NM600) is delivered and followed 9 days later by GD2 *TRAC-*CAR T cells, the residual radiation dose to the CAR T cells is estimated to be below 1 Gy, suggesting a minimal impact on CAR T cell viability ^25^. This may explain why the combination of GD2 *TRAC-*CAR T cells with 1.8 Gy mean tumor dose was more effective than with 3.6 Gy when RPT was administered 9 days before GD2 *TRAC-*CAR T cell infusion.

The interaction between the death receptor Fas on tumor cells and its ligand FasL on T cells is a well-known apoptosis-inducing pathway ^40, 41^. Both EBRT and RPT (^90^Y-based) have been shown to upregulate Fas expression on tumor cells, enhancing immune cell-mediated killing ^42, 43^. Our results suggest that ^177^Lu similarly upregulates Fas on neuroblastoma cells, potentially triggering a CAR-independent mechanism for inducing apoptosis of the tumor. This Fas-mediated pathway may be critical for the eradication of antigen-negative tumor cells by T cells and CAR T cells, addressing a barrier of CAR T cell therapy—antigen escape ^33^.

Single cell proteomics demonstrated enrichment of secreted effector molecules by GD2 *TRAC-* CAR T cells when co-cultured with irradiated neuroblastoma tumors. Simultaneous upregulation of perforin and granzyme with TNFα and IFNγ production indicate CD8^+^ GD2 *TRAC-*CAR T cells co-cultured with GD2+ neuroblastoma cells irradiated by ^177^Lu-based RPT indicates this approach maximally activates CAR T cells ^44^. CAR T cells secreting multiple cytokines have been associated with favorable clinical responses in high-grade lymphoma patients treated with CAR T cells, suggesting that this therapeutic approach could have potential clinical relevance ^45^. While our results demonstrated the feasibility of using RPT to enhance suboptimal CAR T cell function and antitumor activity against the solid tumor neuroblastoma, we did observe reduced efficacy with the solid tumor melanoma, potentially due to its inherent radioresistance compared to neuroblastoma ^46^. Thus, caution should be exercised in extrapolating data across all solid tumor histologies.

Several limitations in our study should be noted. The use of an immunodeficient human xenograft mouse model precluded the evaluation of this therapeutic combination in an immunocompetent system, which could have allowed for observations regarding CAR T cells and other lymphocytes like endogenous T, B and NK cells. Additionally, we did not assess the effects of this combination therapy in a widely metastatic disease model, so it is yet known if the degree of tumor burden influences CAR T cell efficacy.

In summary, our study demonstrates that low dose radiation delivered by ^177^Lu-based RPT enhances the efficacy of GD2 *TRAC-*CAR T cells to eradicate GD2+ solid tumors like neuroblastoma, and to a lesser degree, melanoma, by enhancing antigen-dependent and potentially a CAR-independent mechanisms of cytotoxicity. Complete tumor regression and enhanced overall survival was observed without evidence of end-organ toxicity or xenogeneic GVHD. Factors such as the sequence of administration, radiation dose, radionuclide half-life and linear energy transfer, and timing between RPT and CAR T cell infusion must be carefully considered. Low-dose RPT can enhance otherwise suboptimal CAR T cell function against solid tumors, though additional studies with alternate sources of radiation and CAR T products are required for further optimization in different solid tumor types.

## Acknowledgments

We would like to thank Tracy Berg for editorial review of this manuscript. We also thank the outstanding veterinary staff in the WIMR vivarium for their care of mice throughout *in vivo* studies.

## Contributors

Quaovi H. Sodji: Conceptualization, Resources, data curation, supervision, formal analysis, investigation, methodology, validation, visualization, resources, funding acquisition, writing-original draft, writing-review & editing.

Amanda Shea: investigation, validation, writing-review & editing.

Dan Cappabianca: investigation, validation, writing-review & editing.

Matthew H. Forsberg: formal analysis, investigation, validation, visualization, writing-review & editing.

Jens C. Eickhoff: formal analysis, writing-review & editing.

Malick Bio Idrissou: investigation, validation, writing-review & editing.

Andy S. Ollendorff: investigation, validation, writing-review & editing.

Ohyun Kwon: investigation, validation, writing-review & editing.

Irene M. Ong: formal analysis, writing-review & editing.

Reinier Hernandez: Resources, writing-review & editing.

Jamey Weichert: Resources, writing-review & editing.

Bryan P. Bednarz: Resources, writing-review & editing.

Krishanu Saha: Resources, writing-review & editing.

Paul M. Sondel: Resources, writing-review & editing.

Christian M. Capitini: conceptualization, resources, funding acquisition, writing-review & editing.

Zachary S. Morris: conceptualization, resources, funding acquisition, writing-review & editing.

## Funding

The project was supported by the Clinical and Translational Science Award (CTSA) program, through the NIH National Center for Advancing Translational Sciences (NCATS), grant UL1TR002373. We thank the staff of the University of Wisconsin Carbone Cancer Center (UWCCC) Biostatistics Shared Resource for their valuable contributions to this research. The authors also thank the UWCCC Flow Cytometry core facility and Small Animal Imaging and Radiotherapy core facility. Shared research services at the UWCCC are supported by Cancer Center Support Grant P30 CA014520. We acknowledge generous support from the National Cancer Institute (NCI)/National Institute of Health (NIH) K08 CA285941 (QHS), P01 CA250972 (RH, JW, JCE, IMO, BPB, PMS, ZSM), R01 CA278051 (KS and CMC), S10 S10OD028670 (MILabs microSPECT/CT) University of Wisconsin (UW)-Madison Office of the Vice Chancellor for Research and Graduate Education with funding from the Wisconsin Alumni Research Foundation (RH, JW, KS, PMS, CMC, ZSM), St. Baldrick’s Foundation Empowering Pediatric Immunotherapy for Childhood Cancers (EPICC) Team Grant (KS, PMS, CMC, ZSM), the Midwest Athletes Against Childhood Cancer (MACC) Fund (PMS, CMC, ZSM) and Hyundai Hope on Wheels (KS and CMC). The contents of this article do not necessarily reflect the views or policies of the Department of Health and Human Services, nor does the mention of trade names, commercial products, or organizations imply endorsement by the US Government.

## Conflict of Interest

QHS is an inventor on patent applications related to this publication.

AS: None

DC: None

MHF is an inventor on patent applications related to this publication.

JCE: None

MBI: None

ASO: None

OK: None

IMO: None

RH received patent royalties from Wisconsin Alumni Research Foundation; consulting fees from Archeus Technologies Inc and Monopar Therapeutics.

JW is the founder and member of the Science Advisory Board of Archeus Technologies which holds the licensing rights to NM600 chelates.

BPB received stock/stock options from Voximetry Inc.

KS received grant support from Synthego and Spotlight Therapeutics; patent royalties from Wisconsin Alumni Research Foundation; honoraria from ISCT, is an inventor on patent applications related to this publication; Scientific Advisory Board Member for Notch Therapeutics and Andson Biotech.

PMS received support from the University of Wisconsin, Midwest Athletes For Childhood Cancer and the National Cancer Institute.

CMC received honoraria from Bayer, Nektar Therapeutics, Novartis, WiCell Research Institute, consulting fees and equity interest from Elephas and is an inventor on patent applications related to this publication.

ZSM is a member of the Scientific Advisory Boards for Archeus Technologies, Seneca Therapeutics, Cali Biomedical, and NorthStar Medical Radioisotopes; a consultant for Johnson & Johnson; received royalties from patents held by the Wisconsin Alumni Research Foundation; holds equity options from Archeus Technologies, Cali Biomedical, and Seneca Therapeutics; received research funding provided to the University of Wisconsin from Point Biopharma and Telix Pharmaceuticals and is an inventor on patent applications related to this publication.

## Patient consent for publication

Not applicable

## Ethics approval

T cells were isolated from peripheral blood of healthy donors under a University of Wisconsin-Madison IRB-approved protocol (2017-1070). All procedures were approved by the Institutional Animal Care and Use Committee at the University of Wisconsin-Madison (M005915; M005670; M006815).

## Data availability statement

Data available upon reasonable request

## Supplementary Figures

**Figure S1:**
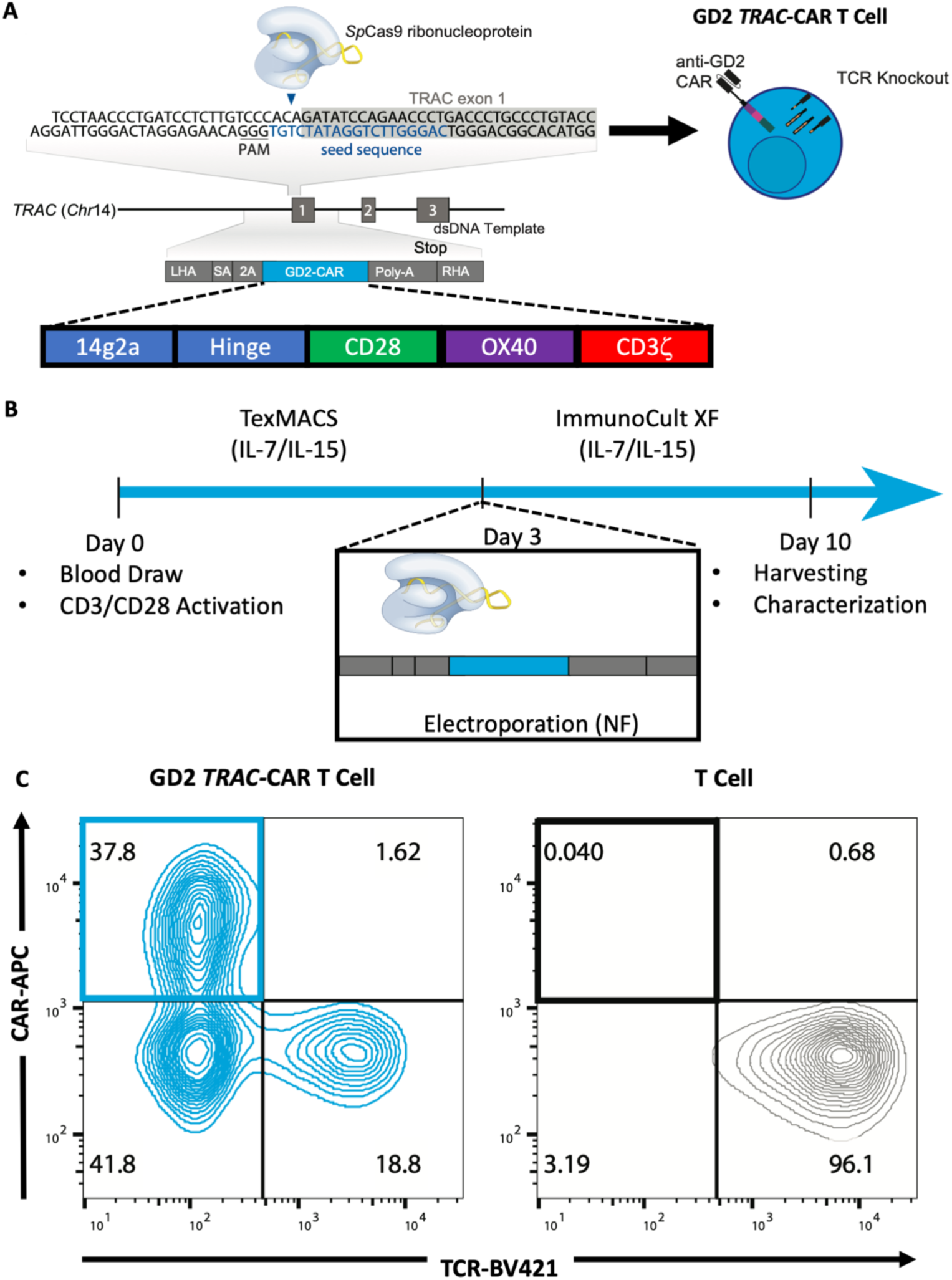
Virus-free GD2 *TRAC*-CAR T production by CRISPR/Cas9 and characterization. **A)** CRISPR/Cas9 knock-in of a third generation GD2 CAR construct into the *TRAC* locus on chromosome (chr) 14. **B)** Conditions and timing of activation, electroporation and expansion of GD2 *TRAC-*CAR T cells**. C)** Characterization by flow cytometry of TCR KO efficiency and GD2 CAR knock-in expression. 2A, cleavage peptide; CAR, chimeric antigen receptor; LHA, left homology arm; NF, Nucleofection; Poly-A, rabbit b-globin polyA terminator; RHA, right homology arm; SA, splice acceptor; TRAC, T cell receptor alpha constant gene; RNP, ribonucleoprotein; PAM, protospacer adjacent motif; TCR, T cell receptor.

**Figure S2:**
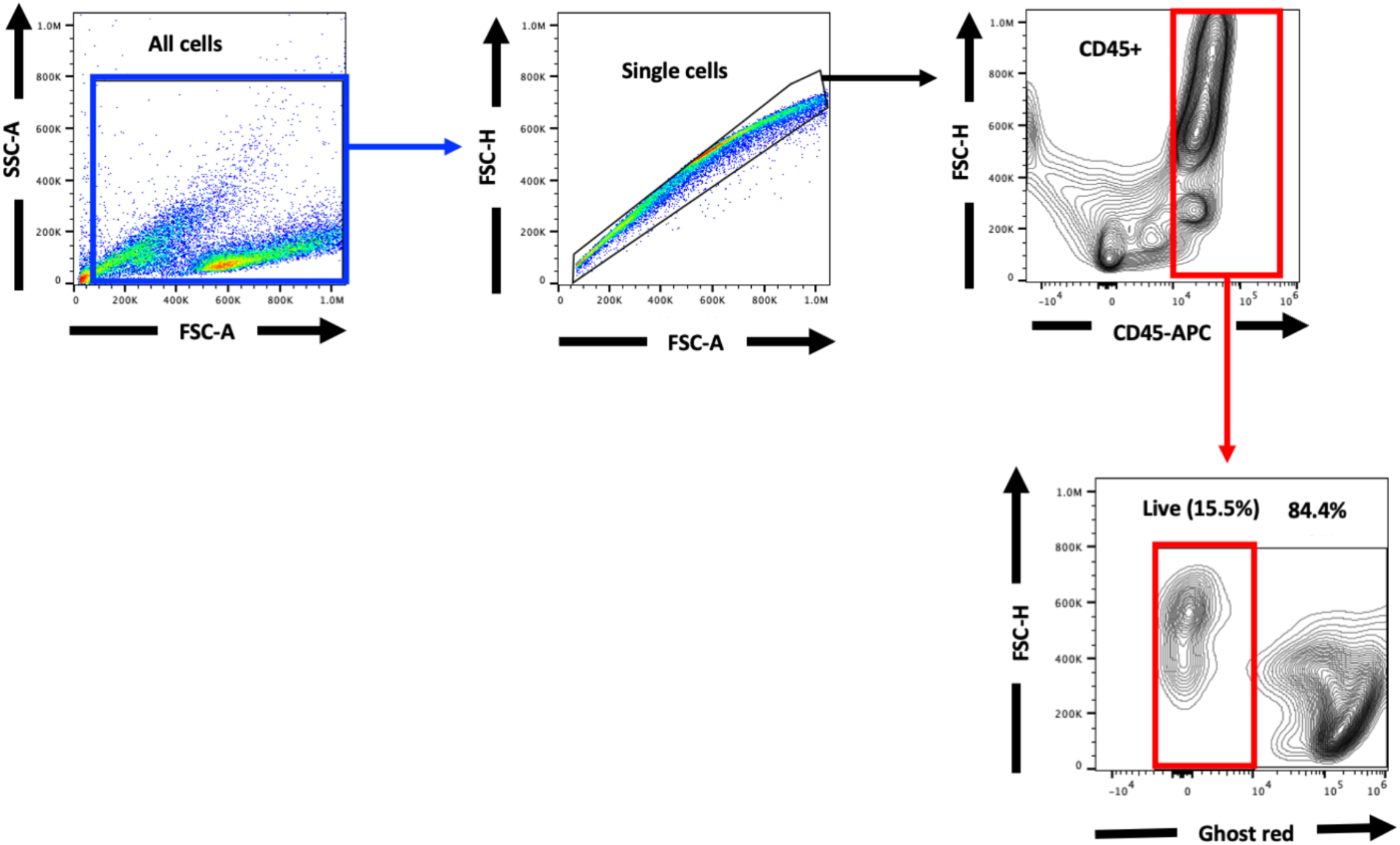
Gating strategy for the detection of *TRAC-*CAR T cells by flow cytometry.

**Figure S3:**
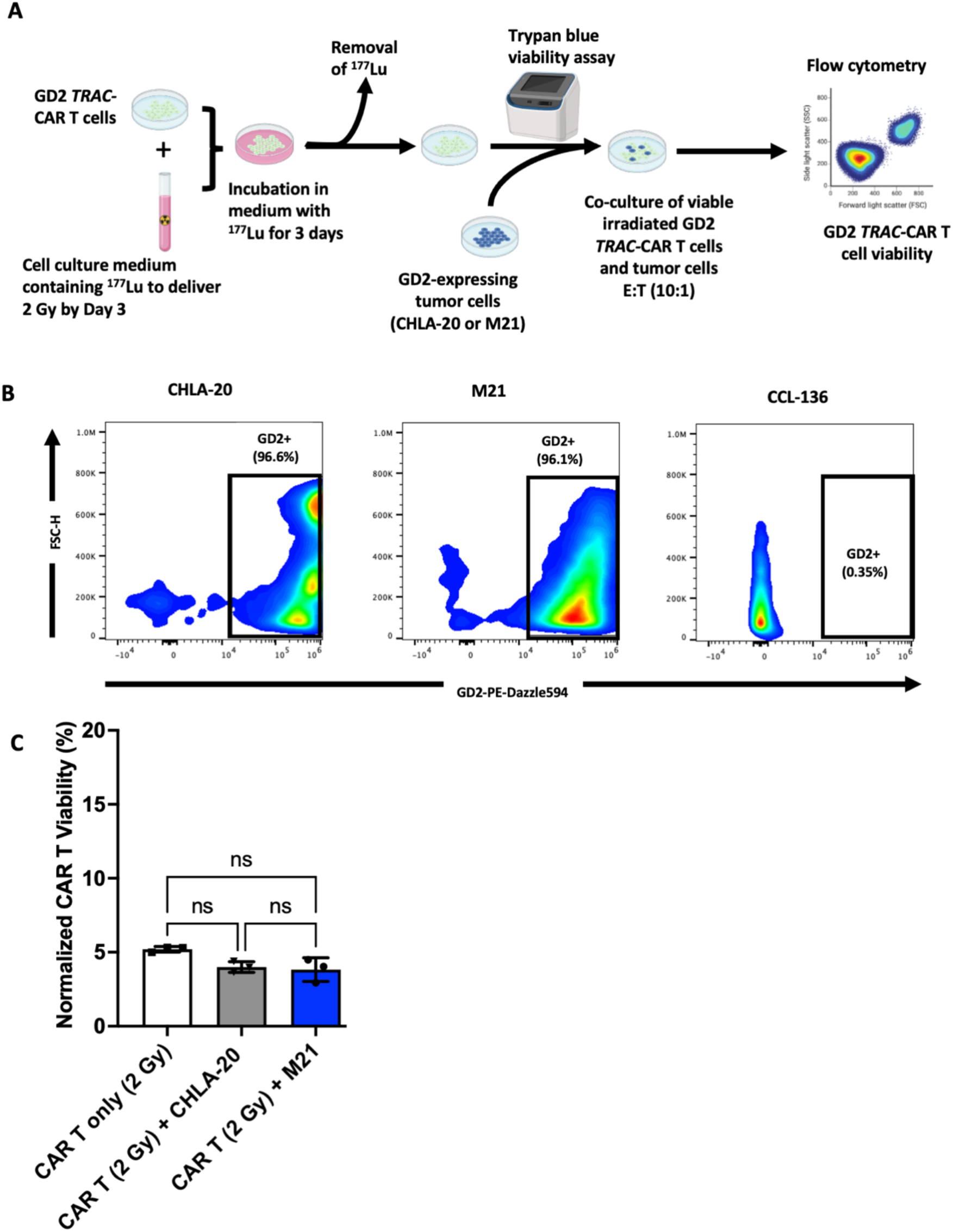
Antigen stimulation after irradiation of GD2 *TRAC-*CAR T cells does not rescue CAR T viability. **A)** Experimental schema. **B)** GD2 expression in CHLA-20 neuroblastoma, M21 melanoma and CCL-136 rhabdomyosarcoma cells. **C)** GD2 *TRAC-*CAR T cell viability following irradiation by ^177^Lu for 3 days and then co-culture with no tumor, CHLA-20 or M21. One-way ANOVA with Tukey’s multiple comparisons. Error bars indicate SD. ns: not significant.

**Figure S4:**
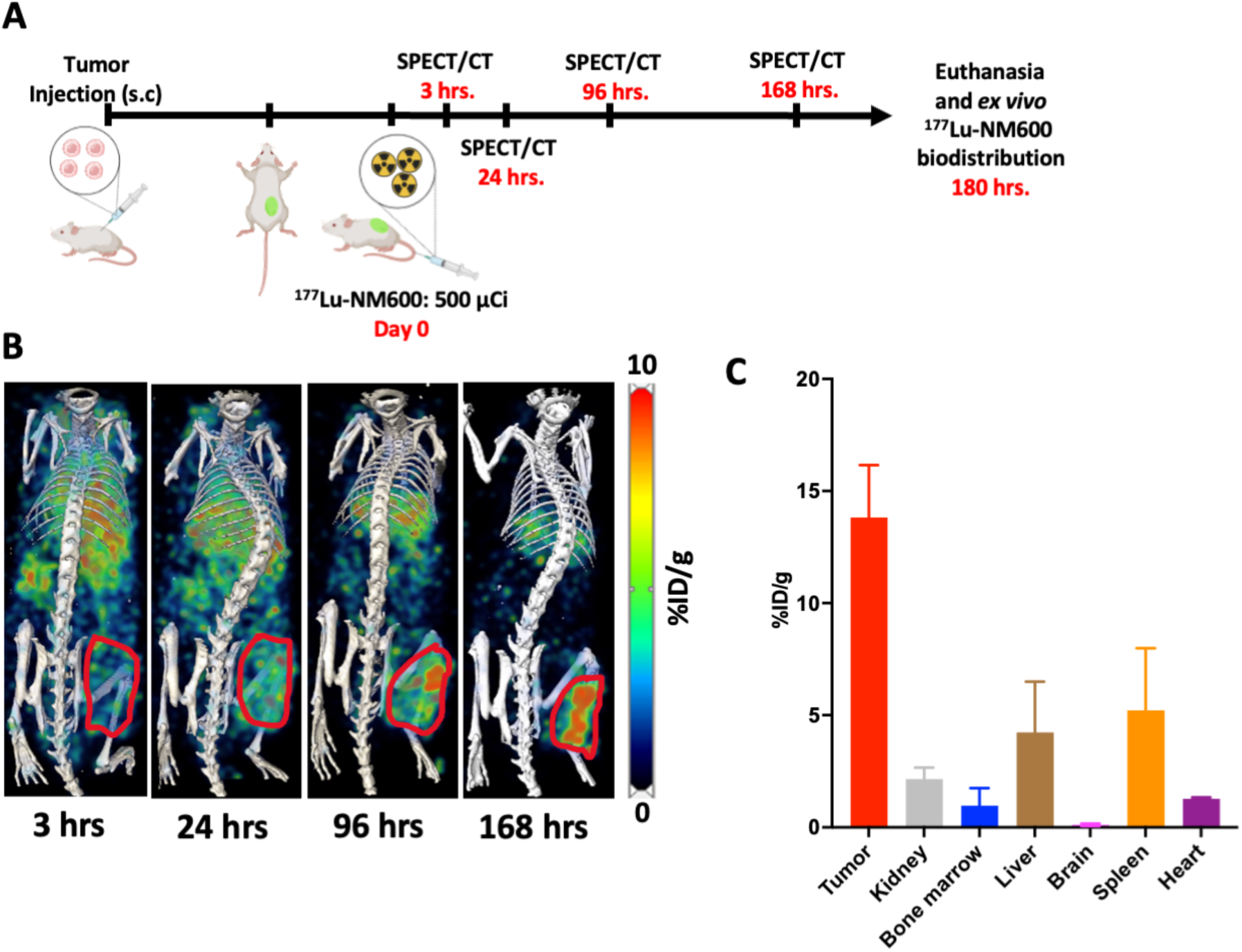
NM600 delivery of ^177^Lu to a human melanoma xenograft model. **A)** Experimental schema. **B)** Representative maximum intensity projection (MIP) SPECT/CT of ^177^Lu-NM600 in an M21 tumor-bearing NRG mouse. **C)** *Ex vivo* biodistribution of ^177^Lu-NM600 in M21 xenograft model 180 hrs after IV injection of ^177^Lu-NM600.

**Figure S5:**
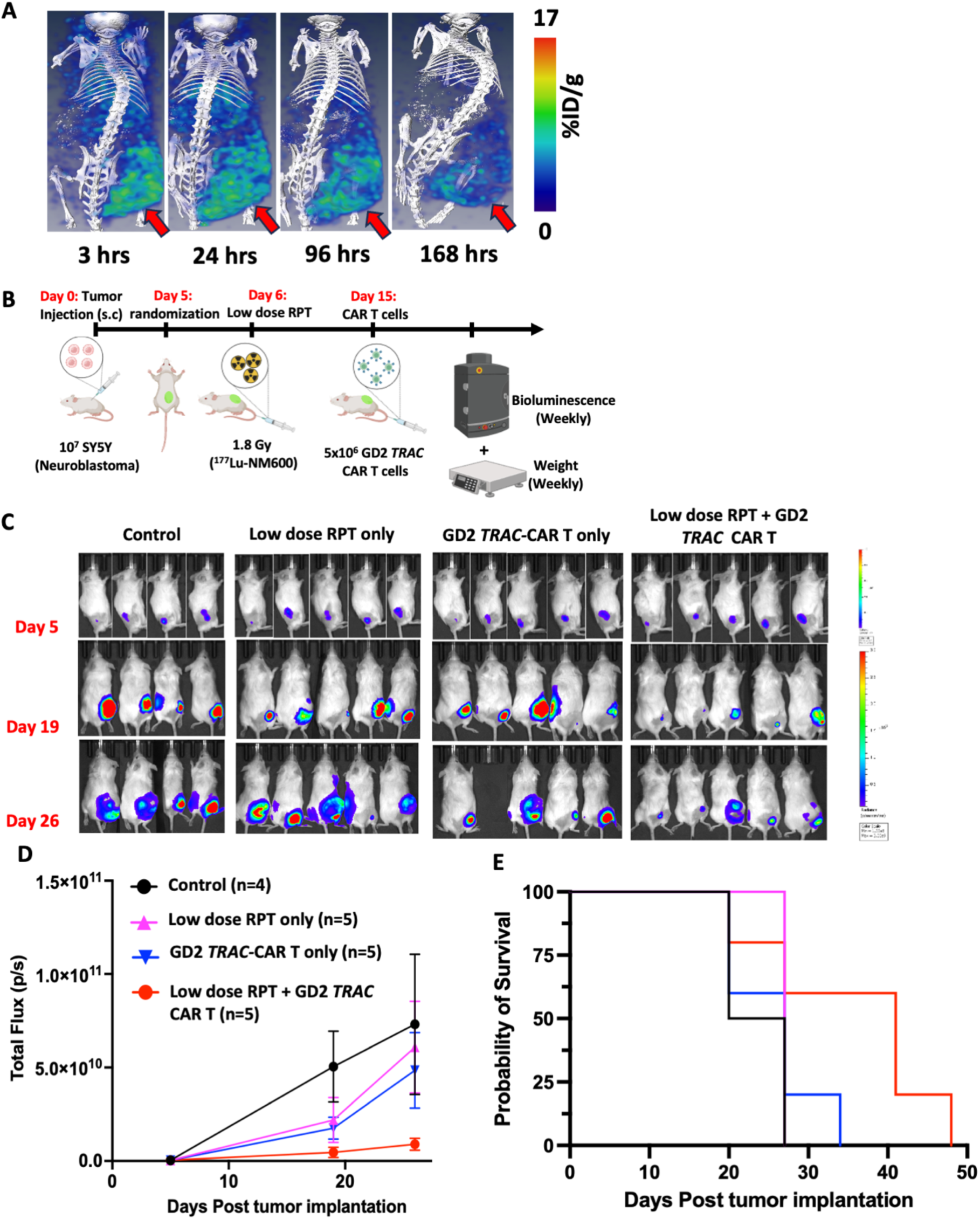
Low mean tumor dose of 1.8 Gy from RPT improves the efficacy of GD2 *TRAC-*CAR T cells in a human xenograft model of neuroblastoma. **A)** Representative maximum intensity projection (MIP) SPEC/CT of ^177^Lu-NM600 in SY5Y tumor-bearing mouse. **B)** Experimental Schema. **C)** Representative bioluminescence images of the mice in all the treatment arms. **D)** Average total flux values of mice in the various treatment arms. **E)** Kaplan-Meier survival curves of all treatment arms.

**Figure S6:**
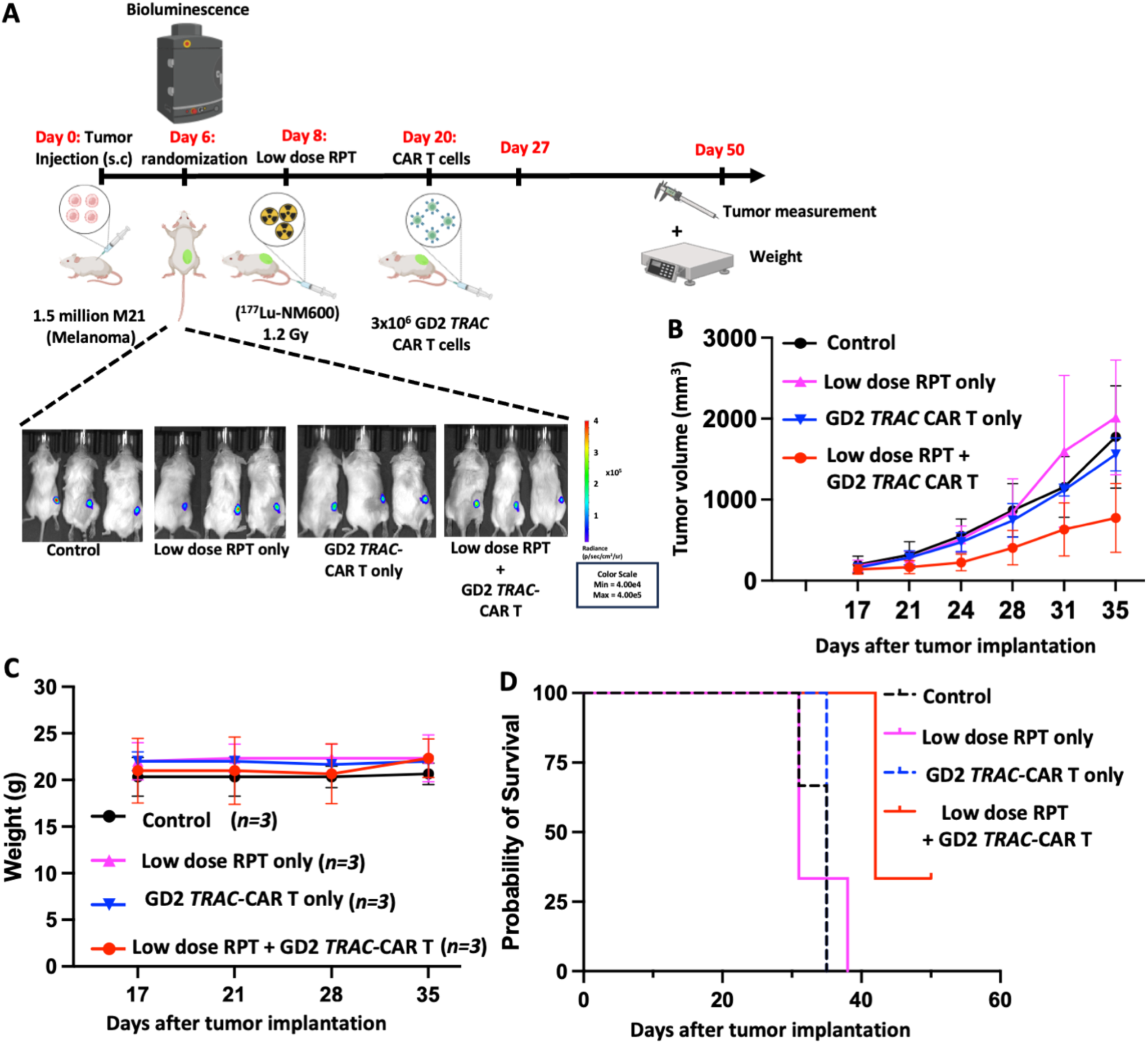
Low mean tumor dose of 1.8 Gy from RPT potentiates GD2 *TRAC-*CAR T cells in a human xenograft model of melanoma. **A)** Experimental Schema. Bioluminescence images of the mice in all the treatment arms. **B)** Tumor growth curves of all treatment arms. **C)** Average weights in grams (g) of all treatment arms over time. **D)** Kaplan-Meier survival curves of all treatment arms.

**Figure S7:**
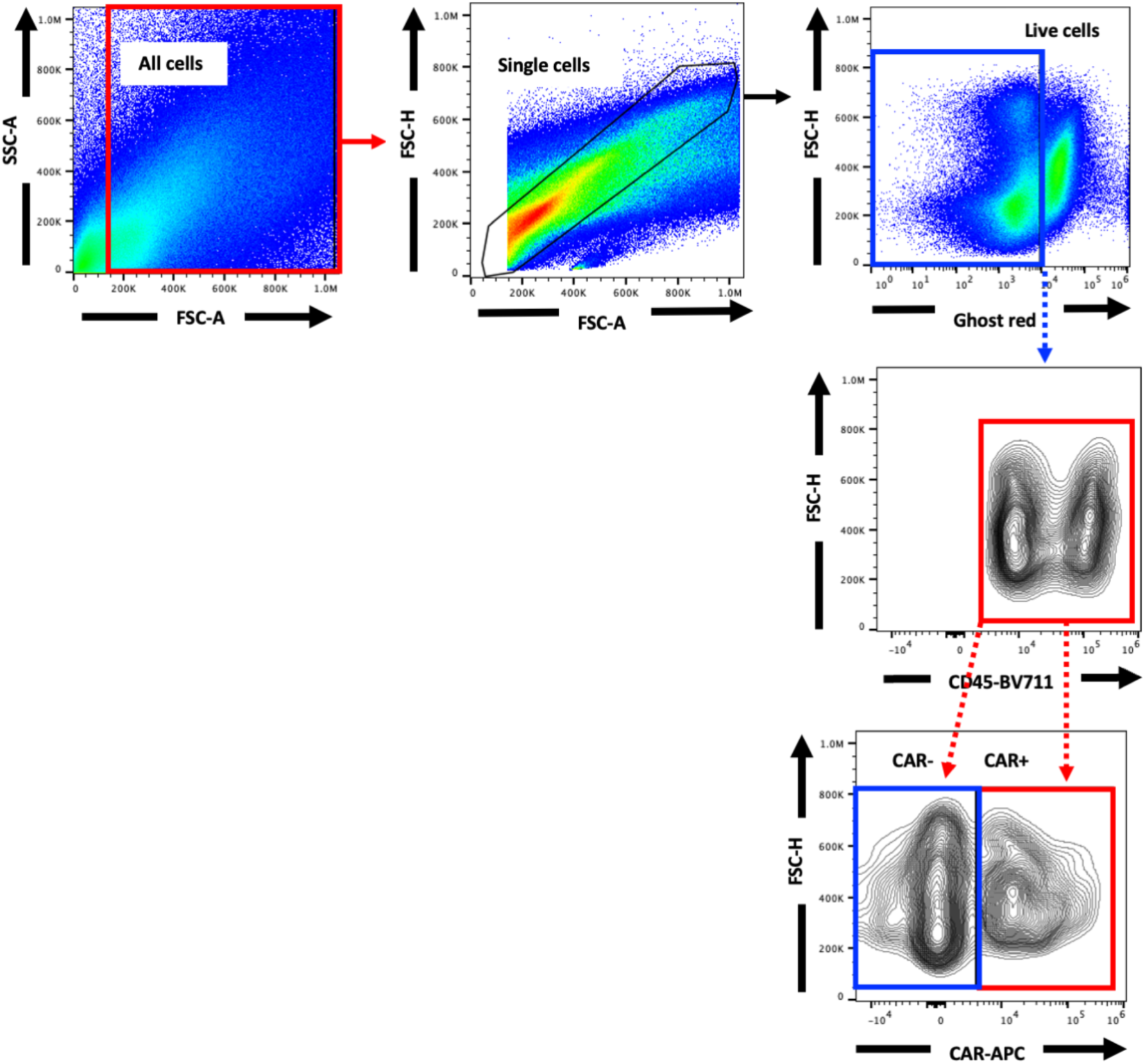
Gating strategy for detecting GD2 *TRAC-*CAR T cells in the TME by flow cytometry.

**Figure S8:**
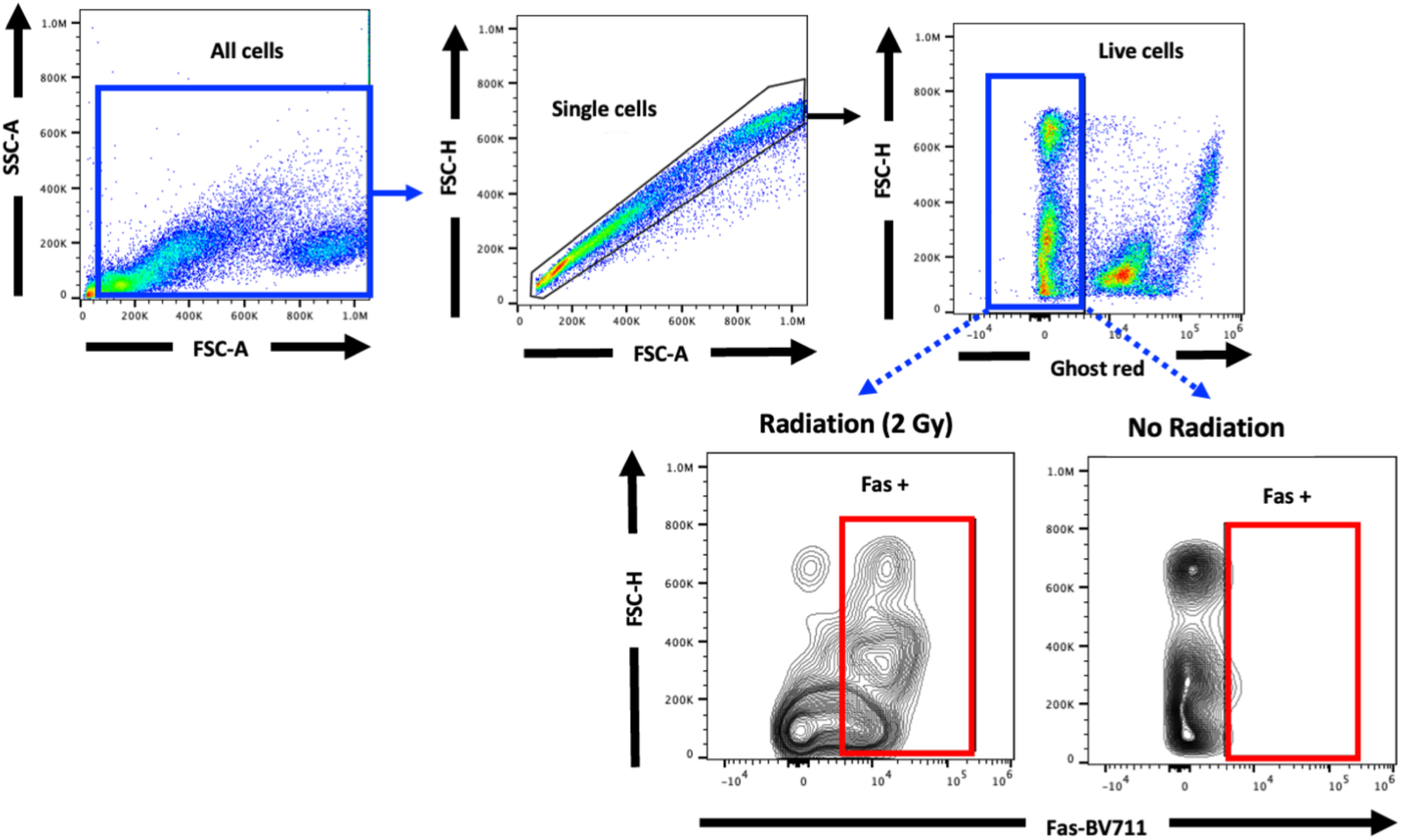
Gating strategy for the detection of Fas expression on irradiated and non-irradiated neuroblastoma cells by flow cytometry.

**Figure S9:**
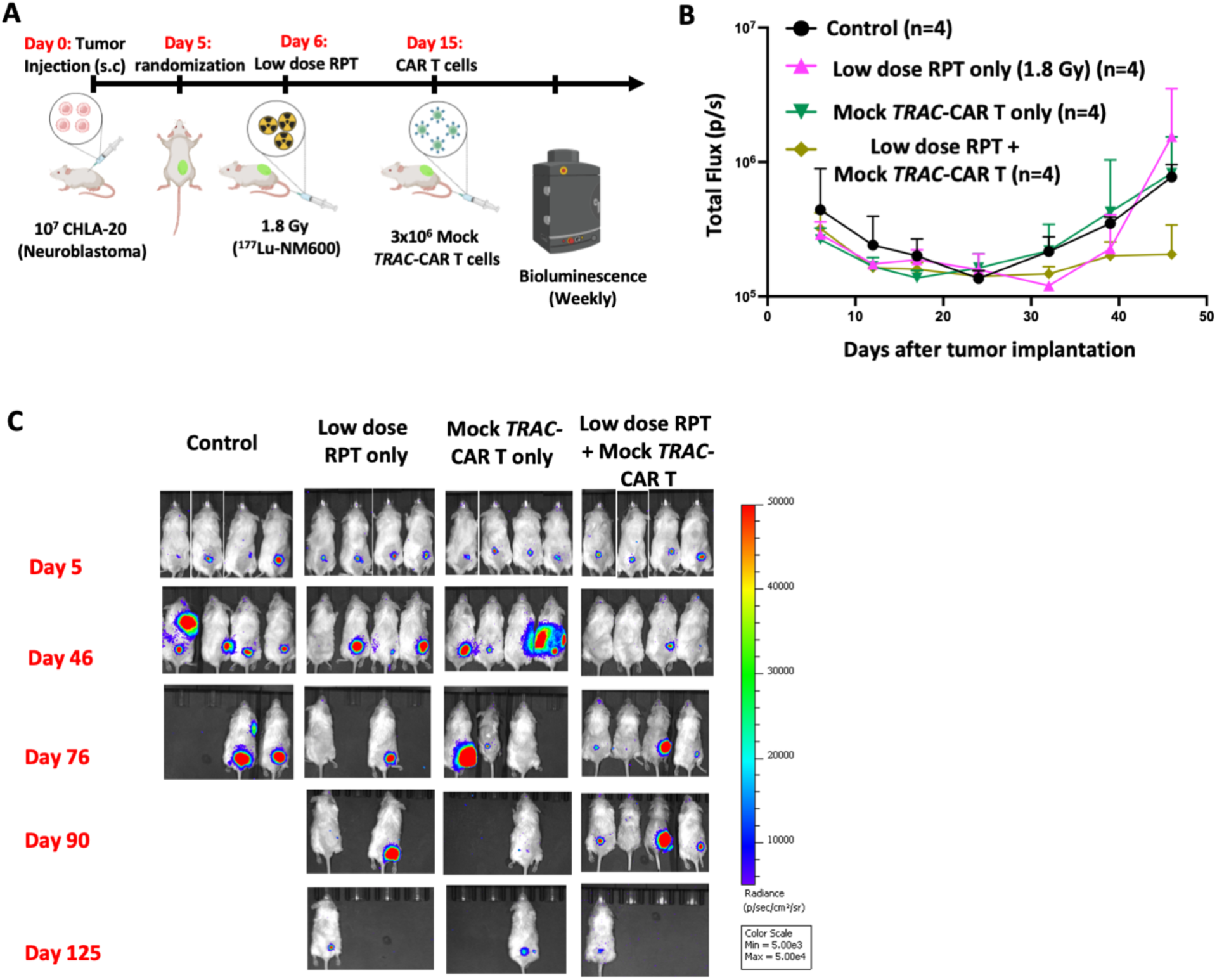
Assessment of antigen independent cytotoxicity treating with low mean tumor dose of 1.8 Gy from RPT followed by mock *TRAC-*CAR T cells. **A)** Experimental Schema. **B)** Total flux values of mice in the various treatment arms. **C)** Bioluminescence images of the mice in all the treatment arms.

